# Axis reset is rate limiting for onset of whole-body regenerative abilities during planarian development

**DOI:** 10.1101/2024.07.19.604337

**Authors:** Clare L.T. Booth, Brian C. Stevens, Clover A. Stubbert, Neil T. Kallgren, Erin L. Davies

## Abstract

Few studies have investigated whether or how regenerative abilities vary across developmental stages of animal life cycles. Determining mechanisms that promote or limit regeneration in certain life cycle stages may pinpoint the most critical factors for successful regeneration and suggest strategies for reverse-engineering regenerative responses in therapeutic settings. In contrast to many mammalian systems, which typically show a loss of regenerative abilities with age, planarian flatworms remain highly regenerative throughout adulthood. The robust reproductive and regenerative capabilities of the planarian *Schmidtea polychroa* (*Spol*) make them an ideal model to determine when and how regeneration competence is established during development. We report that *Spol* gradually acquires whole body regenerative abilities during late embryonic and early juvenile stages. Posterior (tail) regenerative abilities are constitutive, whereas anterior (head) regenerative abilities are dependent on developmental stage, tissue composition of the amputated fragment, and axial position of the cut plane. Stem-like cells are required, but not sufficient, for onset of head regeneration ability. We propose that regulation of main body axis reset, specifically the ability to remake an anterior organizing center, is a rate-limiting factor for establishment of whole-body regeneration competence. Supporting this hypothesis, knock-down of the canonical Wnt pathway effector *Spol β-catenin-1,* a posterior determinant, induces precocious head regeneration under conditions that are normally head regeneration incompetent. Our results suggest that regeneration competence emerges through interactions between cycling stem-like cells, the cellular source of new tissue, and developing adult tissue(s) harboring axial patterning information.

## INTRODUCTION

Regeneration encompasses tissue replacement strategies underlying the maintenance of form and function during homeostasis and the replacement of missing structures following injury. Regenerative abilities are widely distributed across metazoans, from evolutionarily ancient animal clades to vertebrate lineages, and they vary in scope and scale. Some aquatic invertebrates, including hydrozoans, planarians, and acoels, can regrow complete individuals from small tissue fragments^1–3^. Other animals, including many fish and amphibians, can regenerate their appendages such as fins, limbs, and tails^4–6^. Meanwhile, some organisms, including humans and other mammals, have lost these dramatic forms of regenerative ability and can only replace limited tissues and cell types.

Regenerative ability varies not only across species but also across life stages. Most commonly, embryonic and juvenile stages are associated with higher regenerative capacity, and regenerative ability declines during aging^7^. Studies in mice have shown that mammals can regenerate their hearts and digit tips during embryogenesis but this ability declines soon after birth^8–12^. Furthermore, adult stem cell-driven replacement of tissues including blood, muscle, bone, hair follicles, declines during aging^13–17^. Other vertebrates also display losses in competence as they age, such as *Xenopus* frogs which regenerate their limbs as early tadpoles but lose this ability as they approach metamorphosis^18,19^. Within invertebrates, declines in regenerative ability with aging have been reported in groups such as tunicates^20^, insects^21^, and sponges^22^. These age-related declines are likely due to a variety of mechanisms, which include but are not limited to changes in cellular plasticity, stem cell exhaustion, epigenetic changes, and metabolic differences^7,13,23^.

In contrast, an organism gaining regenerative ability over the course of its lifespan may seem counterintuitive. Nonetheless, this phenomenon has been reported in several species, even if the underlying mechanisms have not been thoroughly studied. Juvenile sponges^22^, crinoid larvae^24,25^, and annelid *Capitella teleta* larvae^26^ have all been reported to have more limited regeneration abilities than adults of the same species. The tunicate *Ciona intestinalis*, a close relative of vertebrates, fails to regenerate structures as larvae that adults are capable of regenerating, such as the heart and neural complex^20,27^. Even within vertebrates, some lizard species that are capable of adult tail regeneration do not undergo proper tail regeneration in earlier developmental stages^28^. *Xenopus* frogs also go through a refractory period during tadpole development in which tail regeneration is impaired, but they subsequently improve in regenerative ability^29–31^. However, comparisons of regeneration across life stages have not been thoroughly explored in many highly regenerative model systems.

Planarian flatworms, including *Schmidtea mediterranea (Smed)* and *Dugesia japonica (Djap)*, are well-established models for the study of whole-body regeneration. Small adult tissue fragments excised from any region of the body can produce complete individuals within two weeks^3,32^. In adult *Smed,* regenerative capacity is seemingly inexhaustible in both asexually and sexually reproducing laboratory strains^32,33^. Key to planarians’ regenerative abilities are adult pluripotent stem cells (aPSCs, also called neoblasts), the cellular source of new tissue during homeostasis, asexual reproduction, and regeneration^3,32,34,35^. Cycling somatic aPSCs are widely distributed within the mesenchyme along the main body axis^36–40^. During homeostasis, aPSCs and their progeny interpret patterning information from a constitutively active position control gene network, primarily expressed in the musculature^41^, which influences cell type specification and lineage differentiation^42^, progenitor cell migration, and homing or integration of progenitors into target tissues^35,43–45^. During regeneration, aPSC behavior is influenced by cell-intrinsic and -extrinsic transcriptional responses to injury^46–49^ that facilitate fragment-wide and wound-proximal phases of proliferation^50,51^ and migration of post-mitotic progeny to the wound site to form the regeneration blastema^50,52^.

Blastema establishment and differentiation of missing tissues requires robust axial patterning reset mechanisms. During regeneration initiation, wound-induced expression of key position control genes within the injured muscle, such as *wnt-1* and *notum*^53,54^, and regional patterning information already within the fragment^55,56^ influence main body axis reset. Successful AP axis reset requires the creation of missing signaling center(s), the anterior and posterior poles, which provide inductive signals within the blastema^54,57–62^. Specialized muscle progenitors descended from aPSCs are required to remake the poles^57–59^. Pole re-establishment stabilizes the polarity decision made during regeneration initiation, allowing for recalibration of the position control gene network to produce properly proportioned animals^63,64^.

Studies illuminating whole-body regeneration mechanisms have been conducted in asexually reproducing *Smed* and *Djap* adults, where regenerative abilities are constitutive. Few studies have examined whether or how regenerative abilities vary across the life cycle in planarian species and strains capable of producing directly developing embryos. We employ *Schmidtea polychroa (Spol),* a close relative of *Smed* ^65,66^, to address when regenerative abilities are established during development. We report that the acquisition of whole-body regenerative abilities in *Spol* occurs gradually in late embryonic and early juvenile stages. Intriguingly, differences in regenerative abilities are observed within and across developmental stages. The earliest developmental stages assayed show pronounced axial position-dependent effects on head regeneration ability. Recut assays suggest that headless fragments produced at early developmental stages achieve regeneration competence as their tissues mature.

Next, we sought to determine which system components were required and rate-limiting to establish whole-body regenerative abilities. We show that irradiation-sensitive cells are necessary for regeneration, but their presence is not sufficient for onset of head regeneration ability. We report that the ability to perform AP axis reset is a rate limiting factor for whole-body regeneration competence. Head regeneration-deficient fragments at early developmental stages fail to reform the anterior pole. Perturbation of AP axial information by RNAi knock-down of the canonical Wnt pathway effector *Spol-β-cat-1,* a posterior determinant, induces precocious head regeneration at an early developmental stage under conditions that are normally head regeneration-incompetent. We conclude that establishment of regeneration competence during *Spol* development requires stem-like cells and the maturation of anatomical system(s) that convey axial patterning information necessary for the specification and self-organized assembly of missing structures.

## RESULTS

### *Spol* adults undergo axial position-independent whole-body regeneration

Robust regenerative abilities have previously been reported for *Spol* adults^66^. To determine whether axial cut position impacts regenerative abilities in adult hermaphrodites, we made transverse cuts along the AP axis and monitored fragments for the regeneration of missing tissues at 14 days post-cut (dpc) (**Figure 1A-C**). All fragments were capable of regenerating properly patterned head and tail tissues (**Figure 1A-C**). Midline amputations producing symmetric left (L-) and right (R-) fragments regenerated new tissue along the length of the fragment (**Figure 1D-E**). Parasagittal amputations producing asymmetric tissue fragments, including a thick fragment containing the midline and a thin fragment lacking the midline, both re-established properly patterned regenerates (**Figure 1F-G**). We conclude that *Spol* adults are robust whole-body regenerators, like *Smed*. Head regeneration abilities in *Spol* adults are invariant along the main body axis.

**FIGURE 1:**
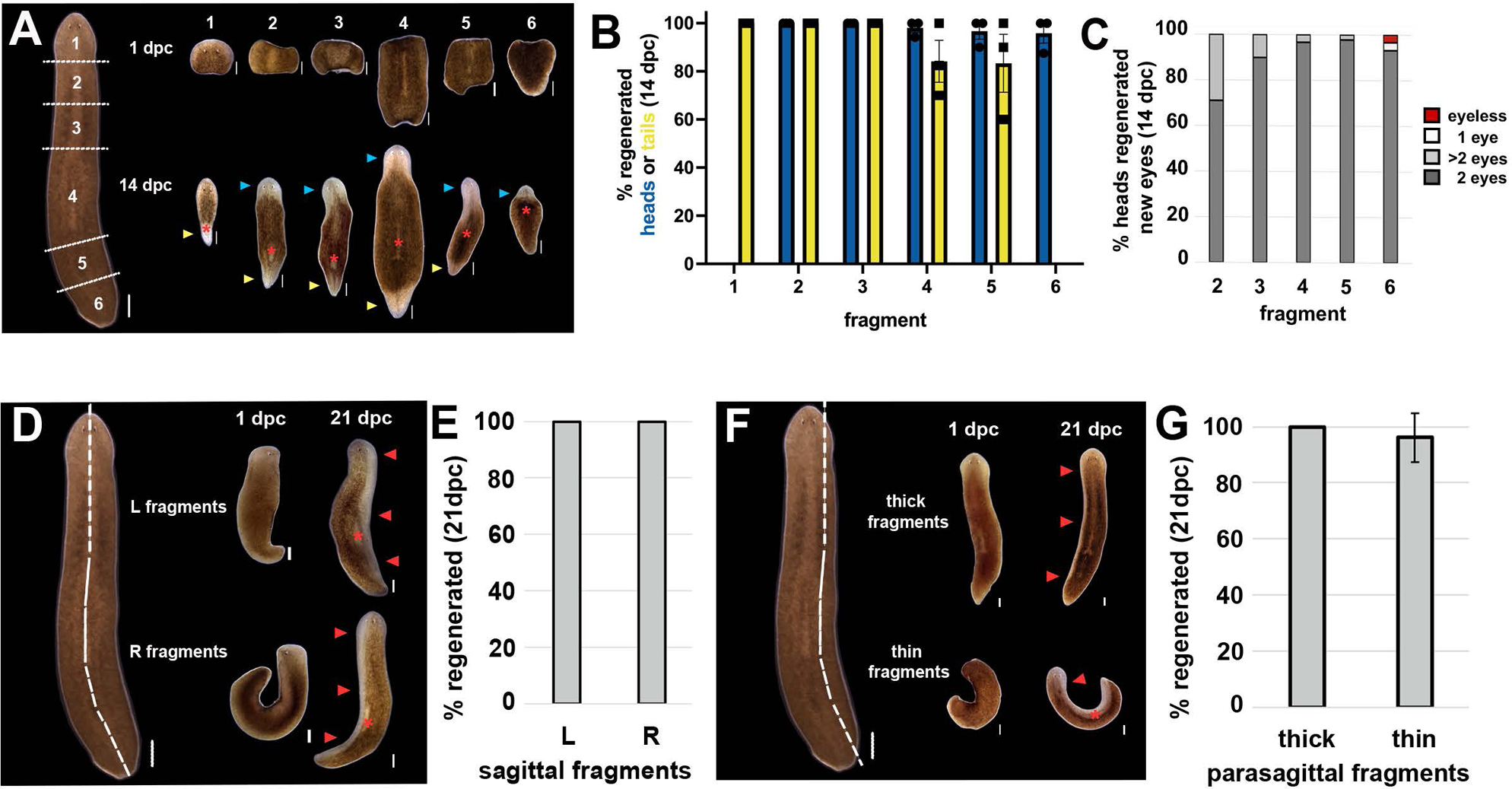
*Spol* adults undergo axial position-independent whole-body regeneration. **1A-C**: **Transverse fragments made along the AP axis regenerate complete animals.** **1A**. Adult hermaphrodites (left) were cut into transverse sections at morphologically defined positions along the anteroposterior (AP) axis (white dashed lines), creating 1: head, 2: anterior prepharyngeal, 3: posterior prepharyngeal, 4: pharyngeal, 5: copulatory, 6: tail fragments. Fragments 1-6 (left to right), top row: 1 day post cut (dpc), bottom row, 14 dpc. Blue arrowheads: regenerated heads. Yellow arrowheads: regenerated tails. Red asterisk: regenerated pharynx. Anterior: up. Dorsal views. Scale bars: Intact adult: 1000 µm. Fragments and regenerates: 500 µm. **1B:** Percent transverse fragments (1-6) that regenerated heads (blue) and tails (yellow) by 14 dpc. **1C**: *Spol* adult fragments produce normally patterned heads at anterior facing wounds. Percent adult transverse fragments 2-6 that regenerated 2 normally patterned eyes (dark gray), >2 eyes (light gray), 1 eye (white), no eyes (maroon) by 14 dpc. **1B-C:** Fragment 1 (n=37), 2 (n=37), 3 (n=35), 4 (n=37), 5 (n=35), 6 (n=36). 3 independent experiments. Error bars: standard error of the mean. **1D-E. *Spol* adults regenerate following sagittal amputation.** **1D**. Adult hermaphrodites (left) were cut along the midline (white dashed line) to create symmetrical left (L-) and right (R-) fragments. Top row: L-fragments. Bottom row: R-fragments. Live images at 1 dpc and 21 dpc. Red asterisk: regenerated pharynx. Red arrowheads: new tissues. **1E**. Percent L- and R- fragments that regenerated missing head, trunk, and tail tissues at 21 dpc. L-fragments (n=27), R-fragments (n=27). 2 independent experiments. Error bars: standard error of the mean. **1F-G. *Spol* adults regenerate following parasagittal amputation.** **1F**. Adult hermaphrodites (left) were cut parasagittally (white dashed line) to create asymmetrical thick and thin fragments. Thick fragments contained the midline and thin fragments lacked the midline. Top row: thick fragments. Bottom row: thin fragments. Live images at 1 dpc and 21 dpc. **1G.** Percent thick and thin fragments that regenerated missing tissues at 21 dpc. Thick fragments (n=30), Thin fragments (n=26). 2 independent experiments. Error bars: standard error of the mean.**1D-F:** Scale bars: Intact *Spol* adult: 1000 µm. Fragments: 500 µm. Red arrowheads: regenerated tissues. Red asterisk: regenerated pharynx. Anterior: up. Dorsal views.

### Whole-body regenerative abilities are acquired gradually during *Spol* development

*Spol* embryos undergo direct development, i.e., they construct the adult body plan during embryogenesis^67^. Like *Smed*, *Spol* embryos are ectolecithal: zygotes are deposited amongst somatically derived yolk cells that are packaged into egg capsules laid by the parents^67^, producing experimentally accessible embryos. We devised a modified *Spol* embryonic staging series consistent with published reports of *Smed* ^73^ and *Spol* ^67,74–77^ development. Construction of the adult anatomy begins during Stage 5 (S5)^67^ (**Figure 2A**). To provide greater temporal and morphological granularity within stages, we subdivided Stage 6 (S6, S6.5) and Stage 7 (S7, S7.5) (**Materials and Methods**). By S7, embryos are elongating along the AP axis and primordia for the brain, eyes, and definitive pharynx are visible (**Figure S1A-B**). Sexually immature juveniles hatch out of egg capsules between 14-21 days post egg capsule deposition (dped) when reared at 20°C. Analysis of spontaneous hatches revealed that 80% of gravid egg capsules hatched between 15-17 dped (**Figure S1C**). We designated the peak of the natural hatching distribution as Juvenile 0 (J0). Formation of the reproductive system occurs post-hatch, during juvenile development.

**FIGURE 2:**
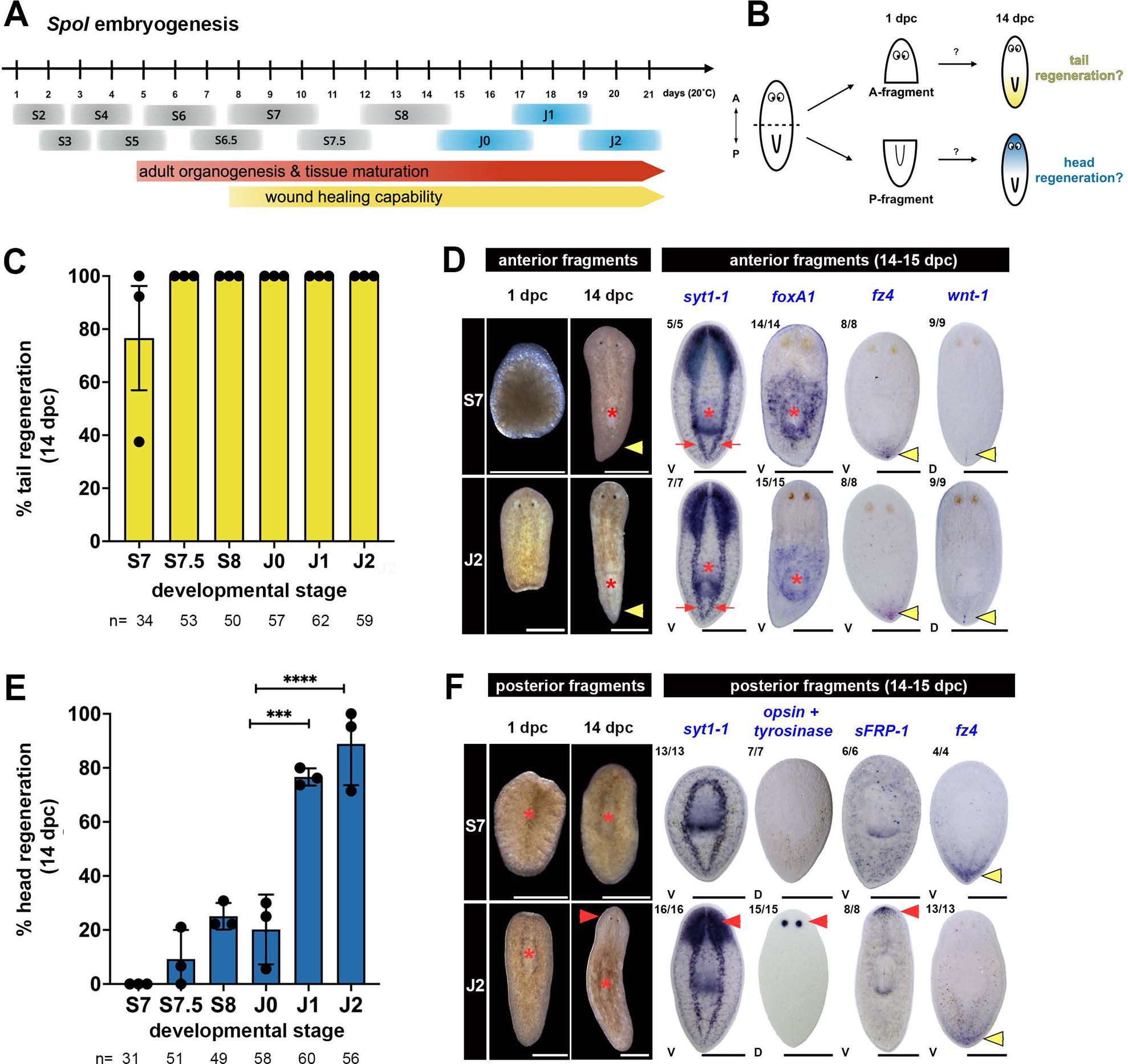
*Spol* acquires whole-body regenerative abilities gradually during late embryonic and early juvenile development. **2A. *Spol* developmental stages**. Embryonic stages (S) S2-S8, Juvenile (J) stages J0-J2. Time: days post-egg capsule deposition (dped), 20°C. Directly developing *Spol* embryos begin constructing the adult body plan during S5. Elongation along the main body axis occurs during S6. Embryo fragments survive transverse amputations from S7 onwards. Spontaneous hatches usually occur during J0 (see **Figure S1C**). Stages of focus in this work: S7-J2. **2B: Embryo dissection scheme.** S7-J2 animals were cut transversely, immediately anterior to the pharynx (dashed line), creating anterior (A-) and posterior (P-) fragments. A- and P- fragments were imaged at 1 and 14 days post-cut (dpc). At 14 dpc, fragments were scored for regeneration of missing tissues and fixed for whole mount in situ hybridization (WISH). **2C-D: Posterior regeneration abilities are constitutive from S7 onwards.** **2C:** Percentage of A-fragments that regenerated missing tails (yellow) at 14 dpc. n: total number of fragments scored over 3 independent experiments. Data points: replicate means. Error bars: standard error of the mean. No statistically significant differences in posterior regenerative abilities were observed for assayed stages (One way ANOVA, (*F*(5,12) = 1.4, *p* = 0.2876)). **2D:** Left: Live images of S7 (top) and J2 (bottom) A-fragments imaged at 1 dpc and 14 dpc. Red asterisk: regenerated pharynx. Yellow arrowhead: regenerated tail. Right: WISH on A-fragment regenerates (14-15 dpc). *syt1-1:* posterior extension of the ventral nerve cords (red arrows). *syt1-1* and *foxA1:* regenerated pharynx (red asterisks)*. fz4:* posterior margin (yellow arrowheads). *wnt-1:* posterior pole (yellow arrowheads). **2E-F: Head regeneration abilities are stage-dependent and increase as animals mature. 2E:** Percentage of P-fragments that regenerated missing heads (blue) at 14 dpc. n: total number of fragments scored over 3 independent experiments. Data points: replicate means. Error bars: standard error of the mean. Statistically significant differences in head regeneration abilities were observed for assayed stages. One way ANOVA, (*F*(5,12) = 44.58, *p*<0.0001). Tukey’s multiple comparison tests: S7 vs J1: padj<0.0001. S7 vs J2: padj<0.0001. S.75 vs J1: padj<0.0001. S7.5 vs J2: padj<0.0001. S8 vs J1: padj<0.0003. S8 vs J2: padj<0.0001. J0 vs J1: padj=0.0001. J0 vs J2: padj<0.0001. **2F:** Left: Live images of S7 (top) and J2 (bottom) P-fragments imaged at 1 dpc and 14 dpc. Red arrowhead: new head. Right: WISH on P-fragment regenerates (14 - 15 dpc). *syt1-1*: new brain (red arrowhead). *opsin-1* and *tyrosinase*: new eyes (red arrowhead). *sFRP-1:* head margin (red arrowhead). *fz4*: posterior margin (yellow arrowhead). Combined scoring data from two independent experiments appears in the upper right corner of each representative image. **2D-F:** Anterior: up. Live images: dorsal views. WISH images: D: dorsal. V: ventral. Scale bars: 500 µm.

To determine developmental onset of regenerative abilities––when embryos were capable of regenerating missing definitive (adult) structures––we performed amputation assays on embryos (S7, S7.5, S8) and early juveniles (J0, J1, J2). Embryos and early juveniles were cut immediately anterior to the developing pharynx, generating anterior (A-) and posterior (P-) fragments (**Figure 2B**). A-fragments contained the head and pre-pharyngeal tissue (**Figure 2D**, 1 dpc), while P-fragments contained the developing pharynx, parapharyngeal region and tail (**Figure 2F**, 1 dpc). At 14 dpc, A-fragments were scored for regeneration of the pharynx, ventral nerve cords, and tail, while P-fragments were scored for regeneration of the head, including the brain and eyes. Amputated S6.5 embryos were deficient at wound closure and lysed after cutting, precluding analysis at earlier developmental stages (data not shown). Most S7 embryos underwent successful wound healing following amputation, albeit a minority of the fragments lysed prior to 14 dpc (**Figure S1D**).

Posterior regeneration abilities were robust and stage independent. A-fragments reliably regenerated new posterior tissues at all stages assayed (**Figure 2C-D**), including the pharynx (**Figure 2D**, *syt1-1* and *foxA1,* red asterisks), a complex organ containing visceral musculature and a peripheral nerve plexus, the ventral nerve cords (**Figure 2D**, *syt1-1,* red arrows), tail tissue (**Figure 2D**, *fz-4,* yellow arrowhead), and the posterior pole (**Figure 2D**, *wnt-1,* yellow arrowhead). A-fragment regenerates exhibited normal gliding locomotion and negative phototaxis (**Movies 1 and 2**).

Strikingly, P-fragments exhibited embryonic stage-dependent differences in head regeneration ability. S7 P-fragments did not regenerate missing anterior tissues (**Figure 2E-F**). S7 P-fragments did not regenerate the bilobed brain, and the ventral nerve cords were fused at the anterior midline (**Figure 2F**, *syt1-1*). S7 P-fragments were eyeless (**Figure 2F**, *opsin + tyrosinase*). S7 P-fragments were headless, but not two-tailed: neither expression of the anterior margin marker, *sFRP-1,* nor expression of the posterior margin marker *fz4*, was observed at anterior-facing wounds in S7 P-fragments at 14-15 dpc (**Figure 2F**, *sFRP-1* and *fz4*). Consistent with the headless phenotype, S7 P-fragments did not exhibit gliding locomotion (**Movie 1**).

S7.5 and S8 P-fragments displayed an intermediate head regeneration phenotype: a minority of them regenerated new head tissue (**Figure 2E, S1E**). S7.5 and S8 P-fragment heads were reminiscent of asexual *Smed* RNAi knock-down animals with small blastemas and compromised anterior regeneration ability (e.g., *prep, foxD, zic-1*)^57–59,78^, and sometimes displayed midline patterning defects (e.g., cyclopia). Head regeneration incidence and quality improved as development proceeded (**Figure 2E-F**). J1 and J2 P-fragments were robust regenerators of new heads containing normally patterned anterior organs, including the brain (**Figure 2F**, *syt1-1*, red arrowhead) and two eyes (**Figure 2F**, *opsin + tyrosinase*, red arrowhead, **S1E**). J1 and J2 P-fragment regenerates also exhibited normal gliding locomotion and negative phototaxis (**Movie 2**).

Next, we sought to disambiguate the contributions of fragment size, tissue composition, and stage to the acquisition of whole-body regeneration abilities. We found no quantitative difference in surface area between regeneration-competent and regeneration-incompetent S7 fragments at 1 dpc (**Figure 3A**), suggesting that tissue composition, rather than fragment size, is the driver of differences in regeneration outcome. Next, we performed transverse amputations at different axial positions in staged embryos (S7, S7.5, S8) and early juveniles (J2) and scored regeneration of missing structures at 14-16 dpc. Posterior regeneration was stage- and axial position-independent (**Figure 3B**). Anterior fragments cut immediately anterior or posterior to the pharynx (prepharyngeal and postpharyngeal cuts, respectively) regenerated missing tail tissues at all stages assayed (**Figure 3B**). In contrast, P-fragments showed marked effects of axial cut position and developmental stage on head regeneration. Posterior fragments were cut immediately posterior to the eyes (**Figure 3 C-D**, P1 fragments), midway between the eyes and pharynx (**Figure 3 E-F**, P2 fragments), or immediately anterior to the pharynx (**Figure 3 G-H**, P3 fragments, **Figure 2B**) and were scored for head regeneration at 14-16 dpc. P1 fragments regenerated properly patterning head tissue at all stages assayed (**Figure 3C-D**, **S2A**). Head regeneration frequency was significantly lower for S7 and S7.5 P2-fragments than S8 and J2 P2-fragments (**Figure 3E-F, S2B**). Consistent with our previous findings, S7 P3-fragments did not regenerate new heads, while S7.5 and S8 P3-fragments displayed some anterior regeneration ability (**Figure 3G-H, S2C**). Regeneration was robust at all axial positions assayed in J2 animals (**Figure 3C-H, S2A-C)**. Finally, we performed sagittal amputations on staged embryos (S7.5, S8) and juveniles (J0, J1, J2) (**Figure 3I**). When survival of Left (L-) and Right (R-) fragments was greater than 50% (**Figure S2D**), we scored regeneration outcomes at 21 dpc. Sagittal regenerates at all stages assayed were capable of regenerating missing head, trunk, and tail tissue by 21 dpc (**Figure 3J-K, S2E**). Together, these data suggest that retention of anterior patterning cues influences onset of head regeneration ability in late-stage *Spol* embryos.

**FIGURE 3:**
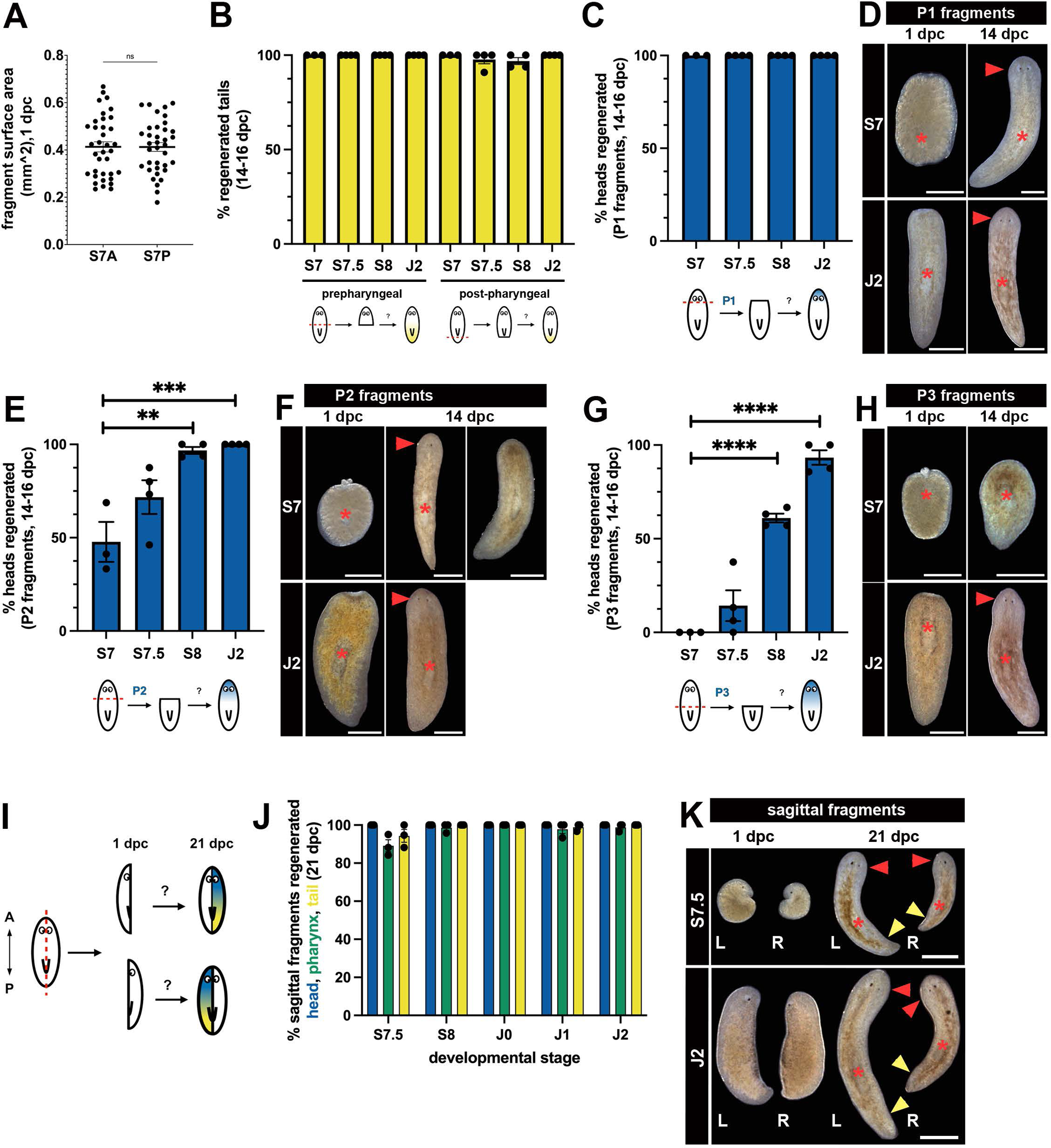
Developmental stage and fragment identity impact head regeneration ability. **3A. S7 A- and S7 P- fragments are similar in size.** No statistically significant difference in the mean surface area (mm^2, horizontal bar) was detected for S7 A- and S7 P-fragment fragments at 1 dpc (Paired t-test, two-tailed, p=0.9757, t=0.03069, df=35). Individual data points are plotted from three independent experiments. Error bars denote standard error of the mean. S7A (n=36), S7 P (n=36). **3B. Posterior regeneration is stage- and axial cut position-independent.** Percent anterior fragments cut prepharyngeally (left) or postpharyngeally (right) from stages S7, S7.5, S8, and J2 that regenerated new tails at 14-16 dpc. Prepharyngeal fragments: S7 (n=46), S7.5 (n=58), S8 (n=63), J2 (n=74). Postpharyngeal fragments: S7 (n=37), S7.5 (n=54), S8 (n=63), J2 (n=77). Error bars: standard error of the mean. S7: 3 independent experiments. S7.5, S8, J2: 4 independent experiments. **3C-H. Head regeneration competence is influenced by developmental stage and cut position.** **3C.** Percent posterior fragments cut immediately posterior to the eyes (cut plane P1) at S7, S7.5, S8, and J2 that regenerated new heads at 14-16 dpc. S7 (n=36), S7.5 (n=48), S8 (n=48), J2 (n=64). Error bars: standard error of the mean. S7: 3 independent experiments. S7.5, S8, J2: 4 independent experiments. **3D.** Live images of S7 (top) and J2 (bottom) P1-fragments imaged at 1 dpc and 14 dpc. **3E.** Percent posterior fragments cut midway between the eyes and pharynx (cut plane P2) at S7, S7.5, S8, and J2 that regenerated new heads at 14-16 dpc. S7 (n=51), S7.5 (n=59), S8 (n=60), J2 (n=69). Error bars: standard error of the mean. S7: 3 independent experiments. S7.5, S8, J2: 4 independent experiments. One way ANOVA, (*F*(3,11) = 13.09, *p*=0.0006). Tukey’s multiple comparison tests: S7 vs S7.5: ns. S7 vs S8: padj=0.0015. S7 vs J2: padj=0.0009. S7.5 vs S8: ns. S7.5 vs J2: padj=0.0360. S8 vs J2: ns **3F**. Live images of S7 (top) and J2 (bottom) P2-fragments imaged at 1 dpc and 14 dpc. **3G.** Percent posterior fragments cut immediately anterior to the pharynx (cut plane P3) at S7, S7.5, S8, and J2 that regenerated new heads at 14-16 dpc. S7 (n=46), S7.5 (n=62), S8 (n=62), J2 (n=77). Error bars: standard error of the mean. S7: 3 independent experiments. S7.5, S8, J2: 4 independent experiments. One way ANOVA, (*F*(3,11) 70.19, *p*<0.0001). Tukey’s multiple comparison tests: S7 vs S7.5: ns. S7 vs S8: padj<0.0001. S7 vs J2: padj<0.0001. S7.5 vs S8: padj=0.0002. S7.5 vs J2: padj<0.0001. S8 vs J2: padj=0.0033. **3H**. Live images of S7 (top) and J2 (bottom) P3-fragments imaged at 1 dpc and 14 dpc. **3I-K. Sagittal amputations produce regeneration competent fragments.** **3I.** Schematic depicting sagittal (midline) amputations performed on late-stage embryos and early juveniles. **3J.** Percent L- and R- sagittal fragments at S7.5, S8, J0, J1, and J2 that regenerated missing head (blue), pharynx (green), and tail (yellow) tissues at 21 dpc. Error bars: standard error of the mean. L+R fragments: S7.5 (n=57), S8 (n=82), J0 (n=92), J1 (n=90), J2 (n=88). 3 independent experiments. **3K.** Live images from S7.5 and J2 L- and R- fragments imaged at 1 dpc and 21 dpc. **3 D, F, H, K**: Anterior: up. Dorsal views. L and R: Left and right sagittal fragments. Red asterisk: pharynx. Red arrowhead: regenerated head. Yellow arrowhead: regenerated tail. Scale bars: 500 µm.

We sought to determine whether regeneration-incompetent fragments could acquire regenerative abilities after maturation by conducting re-amputation experiments. S7 P-fragments (cut at the P3 position) were kept intact or subjected to secondary wounds when they were chronologically equivalent to head regeneration-competent J2 animals (**Figure 4A**). Unsurprisingly, S7 P-fragments that remained intact did not regenerate new heads (**Figure 4B-C**). S7 P-fragments that received injuries associated with minimal tissue loss, such as re-cuts to re-open the initial wound or dorsal incisions above the pharynx, regenerated heads infrequently (**Figure 4B-C**). However, S7 P-fragments that were re-amputated through the pharynx, posterior to the original wound site, produced tail fragments that underwent whole-body regeneration at high frequency (**Figure 4B-C**). Since tail fragments produced through re-amputation were smaller than S7 P-fragments, it is unlikely that S7 P-fragment size falls below a threshold required for head regeneration. Rather, these re-amputation experiments indicate that the composition and/or maturity of the posterior fragments determines head regeneration competence in embryos.

**FIGURE 4:**
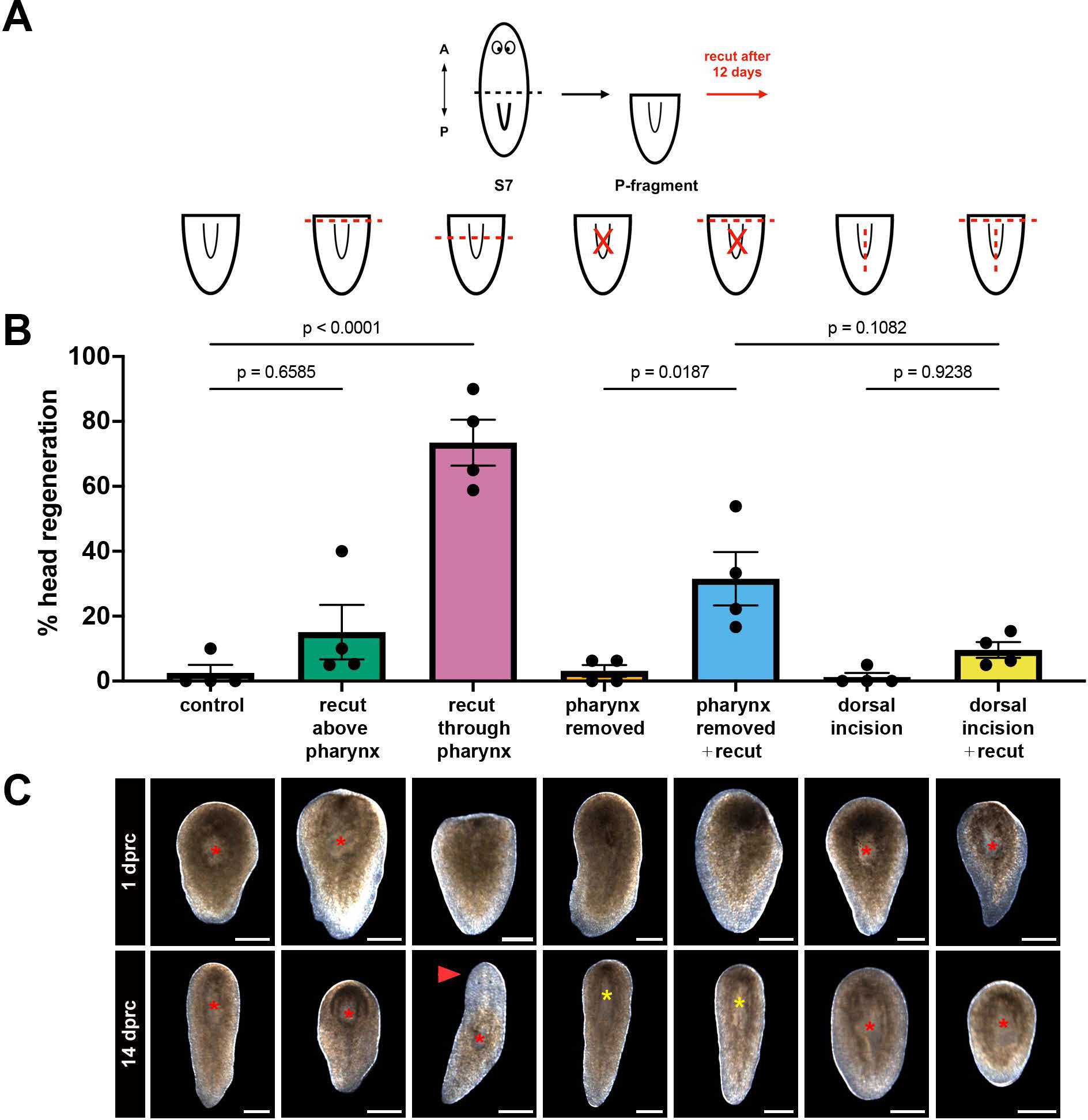
Tail tissue maturation improves head regeneration ability after re-amputation. **4A.** Re-cut experiment schematic. S7 P-fragments were wounded and/or re-cut 12 days after the first amputation and assayed for head regeneration 14 days post-recut (dprc). **4B. Head regeneration frequency after re-cut.** P-fragments re-cut above the pharynx did not show a significant increase in regeneration frequency while those re-cut through the pharynx did. Pharynx removal and dorsal incision alone did not trigger head regeneration, though pharynx removal and re-cut modestly increased regeneration frequency compared to pharynx removal alone. Four replicates were performed for each re-cut paradigm (total n = 63, 64, 61, 61, 61, 66, 65). Statistics computed using Tukey’s multiple comparisons test. **4C**. Representative images of re-cut fragments 1 day post re-cut (dprc) and 14 dprc. A pre-existing (red asterisks) or regenerated (yellow asterisks) pharynx is present in all fragments. Fragments re-cut through the pharynx regenerate heads (red arrowhead) with high frequency. Scale bars = 250 µm.

Differences in the observed head regeneration frequencies dependent on re-cut position may be attributable to the amount or type of tissue removed. To test whether presence of the pharynx influenced head regeneration frequency after re-cut, we surgically removed the pharynx through a dorsal incision. As a control, we performed the same dorsal incision without pharynx removal. While the pharynx itself regenerated after surgical removal (**Figure 4C**, yellow asterisks), neither pharynx removal nor the dorsal incision alone resulted in improved head regeneration (**Figure 4B-C**). Furthermore, we did not observe a significant difference in regeneration frequency in pharynx-removed fragments that were re-cut compared to dorsal-incision fragments that were re-cut at the initial wound site (**Figure 4B**, p=0.1082, Tukey’s multiple comparisons test). These results suggest that pharynx removal does not have a significant impact on regeneration frequency after re-cut. Juxtaposition of the remaining pharyngeal pouch and the wound site may impede regeneration by impairing the movement and/or accumulation of progenitor cells required to build a blastema, or through the elevated expression of posterior patterning genes such as *wnt11-4, wnt11-5* and *wnt11-6* in the pouch and esophagus^63^. Our results suggest that in juvenile *Spol*, significant tissue removal may be required to trigger head regeneration.

Altogether, our results show that regeneration competence is acquired gradually during *Spol* development, with axial-independent, whole-body regenerative abilities arising shortly after hatching (J1-J2). Tail regeneration is constitutive and position-independent by S7, while pronounced stage- and axial position-dependent effects on head regeneration ability were observed prior to J1. Our usage of the term “regeneration-incompetent” describes an anterior regeneration defect, rather than a complete regeneration block. Severity of head regeneration defects correlated with instances where resetting the AP axis was required. Although fragment size can limit regeneration competence^49^, it is unlikely to cause the head regeneration defects that we observed. Instead, tissue composition, axial position, and developmental stage of the fragment are likely to be critical determinants of head regeneration competence.

### Cycling *piwi-1+* cells are required but not sufficient for whole-body regeneration

Cells expressing *piwi-1*^73^ and other germline multipotency genes (e.g. *Spol-tud-1*^79^ and *Spol-vlg-A*^80^) are present throughout planarian embryogenesis and give rise to progenitor cells necessary for the construction of the adult anatomy. In S7, mesenchymal *piwi-1+* cells are widely distributed along the AP axis (**Figure S3A**). We developed a high dose gamma irradiation protocol to completely and irreversibly eliminate *piwi-1+* cells and their descendants in developing embryos. Treatment of 7 dped egg capsules (estimated stage: S6 - S6.5) with 10,000 Rad was lethal, although most embryos continued to elongate and develop in the short-term (**Figure 5A-C**). Irradiated embryos developed edema by 7 dpi (**Figure 5C**), and ultimately succumbed to tissue lesioning and lysis within 10-20 dpi (**Figure 5B-C**). We confirmed that *piwi-1+* cells and irradiation-sensitive epidermal progenitors marked by *prog-1* and *AGAT-1* were greatly reduced by 3 dpi and effectively depleted within 7 dpi (**Figure 5D**), suggesting that high dose irradiation disrupted tissue development and homeostasis. In contrast, we observed robust and persistent expression of *piwi-1*, *prog-1,* and *AGAT-1* in unirradiated controls (**Figure 5D**).

**FIGURE 5:**
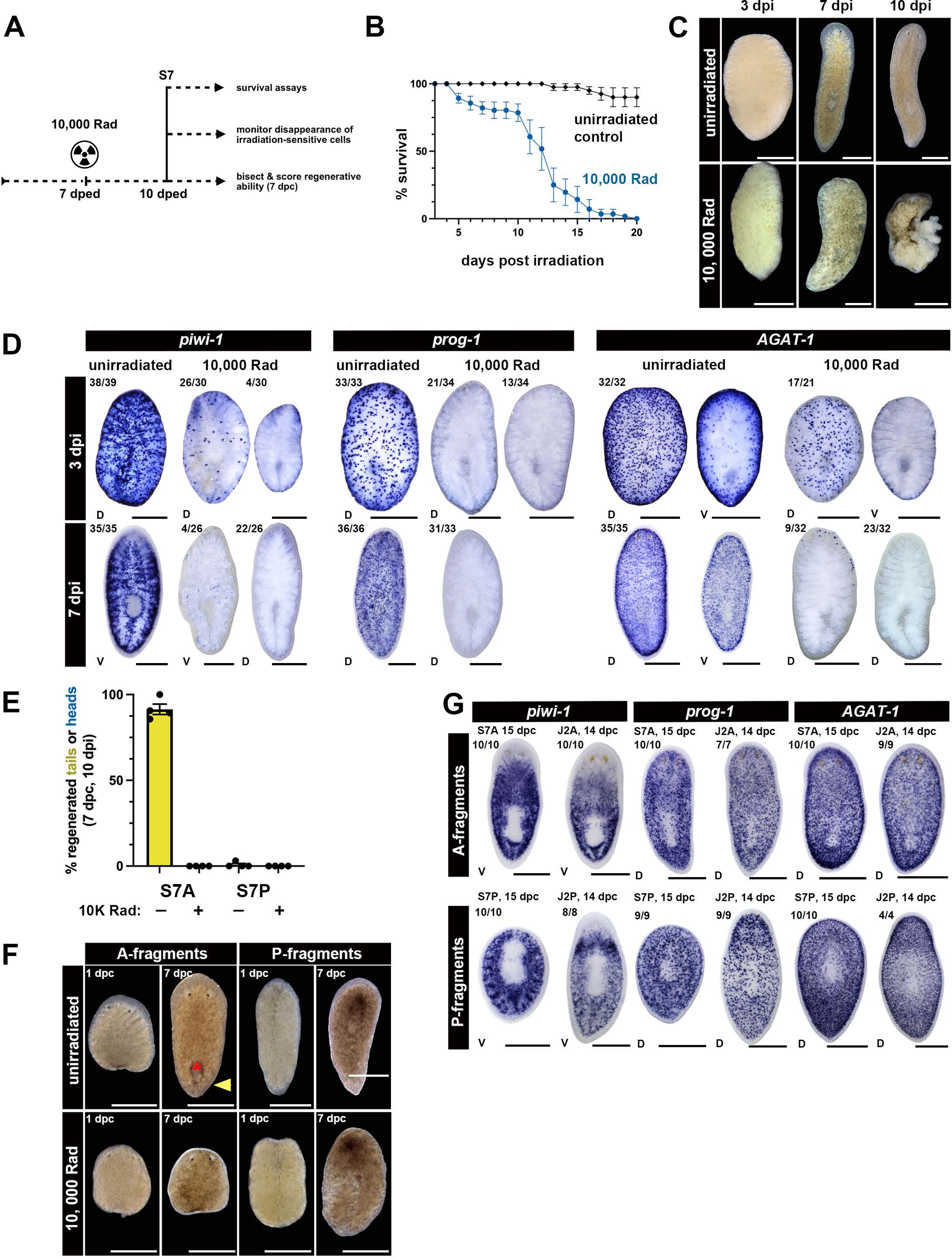
Cycling *piwi-1+* cells are necessary but not sufficient for regeneration competence. **5A-E: Irradiation-sensitive *piwi-1+* cells are necessary for the regeneration of new tissues during development.** **5A.** 7 dped egg capsules were split into unirradiated control and irradiated groups. Irradiated egg capsules received 10,000 Rad. At 3 days post-irradiation (dpi) (10 dped), S7 embryos were harvested from unirradiated and irradiated egg capsules to: 1) monitor long-term survival and onset of irradiation-induced phenotypes, 2) assess the kinetics of *piwi-1+* and differentiating epidermal progenitor cell (*prog-1+* and *AGAT-1+*) loss post-irradiation, and 3) determine whether irradiation-sensitive cells were required for regeneration in S7 embryos. **5B-C. 10,000 Rad treatment at 7 dped is lethal within 10-20 dpi.** **5B**. Percent surviving unirradiated control (black) or 10,000 Rad treated (blue) embryos as a function of time (dpi). Four independent experiments. Unirradiated controls: n=40. 10,000 Rad treated embryos: n=56. **5C.** Live images of unirradiated (top) and 10,000 Rad treated (bottom) embryos dissected at S7(10 dped, 3 dpi) and monitored for one week (17 dped, 10 dpi). By 7 dpi, 82.5 +/− 15.5% irradiated embryos had edema. Bloated embryos eventually burst, leading to tissue loss and lysis within 10-20 dpi. **5D. 10,000 Rad treatment at 7 dped strongly reduces *piwi-1+, prog-1+,* and *AGAT-1+* cell numbers within 3-7 dpi.** WISH with *piwi-1, prog-1,* or *AGAT-1* riboprobes on unirradiated control (left) and 10,000 Rad treated embryos (right) at 3 dpi (top) and 7 dpi (bottom). Combined scoring data from four independent experiments appears in the upper left corner of each representative image. **5E-F. Irradiation-sensitive cells are required but not sufficient for regeneration.** **5E.** Percent unirradiated or 10,000 Rad treated S7 A-fragments that regenerated missing tails (yellow) and S7 P-fragments that regenerated missing heads (blue) at 7 dpc (10 dpi). Total number of fragments scored over 4 independent experiments: unirradiated S7 A n=132, irradiated S7A: n=83, unirradiated S7 P: n=131, irradiated S7 P: n=67. Data points: replicate means. Error bars: standard error of the mean. **5F**. Live images of unirradiated (top) and 10,000 Rad-treated (bottom) S7A- and S7 P-fragments at 1 dpc (4 dpi) and 7 dpc (10 dpi). Red asterisk: regenerated pharynx. Yellow arrowhead: regenerated tail. **5G. *piwi-1+*, *prog-1+*, and *AGAT-1+* cells persist in headless S7 P-fragments.** WISH with *piwi-1*, *prog-1,* or *AGAT-1* riboprobes on unirradiated S7 A- and J2 A- regenerates (top), and S7 P- and J2 P-regenerates (bottom) at 14-15 dpc. Scoring data appears in the upper left corner of each representative image. **5C-G:** Anterior: up. Live images: dorsal views. WISH images: D: dorsal. V: ventral. Scale bars: 500 µm.

To determine whether *piwi-1+* cells are required for regeneration in developing embryos, we performed amputation assays on stage-matched unirradiated control and 10,000 Rad-treated S7 embryos depleted of *piwi-1+* cells at 3 days post-irradiation (dpi). Poor survival of irradiated S7 A- and S7 P- fragments relative to unirradiated controls precluded scoring regeneration later than 7 dpc (**Figure S3B**). A complete regeneration block was observed in irradiated S7 embryos (**Figure 5E-F**), demonstrating that irradiation-sensitive cells are required for regeneration of new tissues during development. Irradiated S7 A-fragments did not regenerate missing posterior structures, while unirradiated S7 A-fragments reliably created new pharynges and tails by 7 dpc (10 dpi) (**Figure 5E-F**). Neither irradiated S7 P-fragments, nor their unirradiated counterparts, produced new head structures (**Figure 5E-F**). We determined that *piwi-1+* cells remain in S7 P- fragments at 14 dpc (**Figure 5G**), and we detected mitotic cells in S7 and J2 A- and P-fragments at 48 hpc (**Figure S3C**). These results indicate that neither *piwi-1*+ cell loss nor a cell proliferation block underlie the head regeneration-incompetent phenotype observed in S7 P-fragments.

To determine whether *piwi-1+* cells in head regeneration-incompetent fragments could produce differentiating progeny, we assayed for persistence of *prog-1+* and *AGAT-1+* epidermal progenitors in unirradiated S7 A- and S7 P-fragment regenerates at 14 dpc. High dose irradiation (**Figure 5D**) or knock-downs that abrogate the production of differentiating epidermal progenitors in *Smed* asexual adults (e.g, *Smed-zfp-1, Smed-p53* RNAi, *Smed-mex3* RNAi) elicit loss of the epidermal lineage^52,81–83^. We observed body-wide distributions of mesenchymal *prog-1*+ and *AGAT-1+* cells in unirradiated S7 A-, S7 P-, J2 A-, and J2 P-fragments at 14 dpc (**Figure 5G**). The persistence of *prog-1+* and *AGAT-1+* cells in headless S7 P-fragments suggests that production of differentiating epidermal lineage cells continues even though head regeneration failed. We hypothesize that *piwi-1+* cells in S7 P-fragments sustain production of lineages necessary for the development and maintenance of “pre-existing” posterior structures.

Our results suggest that gross changes in the number or distribution of *piwi-1+* cells, cycling cell activity, or differentiation potential of *piwi-1+* cell descendants are unlikely to explain the observed stage- and axial position-dependent differences in head regeneration ability. We conclude that irradiation-sensitive cells are required, but their presence is not sufficient, for regeneration in late-stage *Spol* embryos.

### AP axis reset is rate limiting for establishment of whole-body regeneration

AP axis reset is a critical precursor to head regeneration. In adult *Smed*, the anterior pole head organizing center is re-formed in the blastema by 48-72 hpc (reviewed in ^84^). The anterior pole expresses modulators of signaling pathways that control AP axis identity, including inhibitors of Wnt and Activin signaling (*notum* and *follistatin*, respectively^54,85^). When pole formation is disrupted via knockdown of specifier genes including *zic1* and *foxD*, animals fail to regenerate a head, a phenotype reminiscent of headless S7 P-fragments^57–59^. We assayed S7 and J2 P-fragment regenerates for expression of anterior pole markers (*notum, follistatin, foxD)* and head margin markers (*sFRP-1)* at 3 dpc and 14 dpc (**Figure 6A**). J2 P-fragments initiate formation of the anterior pole within 3 dpc, as judged by the appearance of *notum+*, *follistatin*+, and *foxD+* puncta at the blastema midline (**Figure 6A**, red arrowheads). *sFRP-1* expression was also detected around the developing head margin by 3 dpc in regenerating J2 P-fragments (**Figure 6A**). In contrast, S7 P-fragments failed to re-establish the anterior pole, as judged by the absence of *notum+*, *follistatin*+, and *fox-D+* cells at 3 dpc and 14 dpc (**Figure 6A**). *sFRP-1* expression was not detected in S7 P-fragments (**Figure 6A**), further indicating that the blastema did not adopt anterior identity.

**FIGURE 6:**
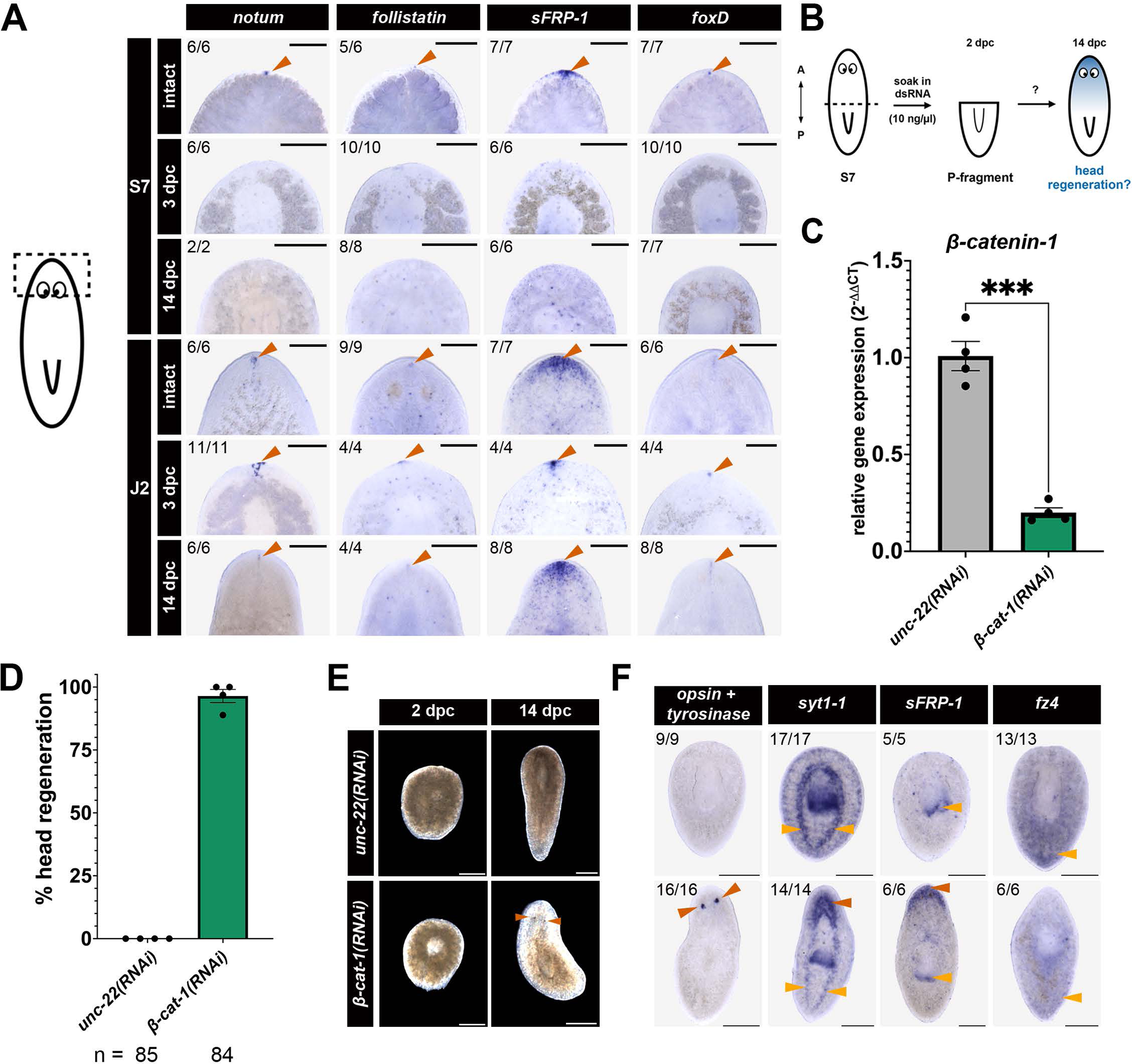
AP axis reset is rate limiting for establishment of head regeneration ability. **6A. The anterior pole is not re-established in headless fragments.** WISH on intact S7 and J2 animals, and S7 and J2 P-fragments at 3 dpc and 14 dpc, with riboprobes that mark the anterior pole (*notum*; *follistatin*; *foxD*, red arrowhead) and the anterior margin and pole (*sFRP-1,* red arrowhead). Scoring data appears in the upper left corner of each representative image. **6B-F**. ***β-cat-1* knock-down elicits precocious head regeneration in S7P fragments.** **6B.** Experimental schematic for knockdowns. S7 animals were bisected and soaked in 10 ng/ul *unc-22* (control) or *β-cat-1* dsRNA for two days. Fragments were scored for regeneration at 14 dpc. **6C.** *β-cat-1(RNAi)* P-fragments at 14 dpc have a significant (p=0.0009, paired t-test) decrease in *β-cat-1* mRNA levels compared to *unc-22(RNAi)* P-fragments at 14 dpc, as measured by RT-qPCR. *β-cat-1* expression is normalized to the expression of the housekeeping gene *ef2*. **6D.** Percent of 14 dpc S7P *unc-22(RNAi)* and *β-cat-1(RNAi)* tail fragments that regenerated heads. **6E.** Representative live images of *unc-22(RNAi)* and *β-cat-1(RNAi)* S7 P-fragments at 2 dpc and 14 dpc. *β-cat-1(RNAi)* tail fragments regenerate heads with eyes (red arrowheads). Scale bars = 250 µm. **6C-E:** RNAi construct information: see Figure S4A. Results shown for four independent experiments. Error bars: standard error of the mean. **6F. WISH with anterior biomarkers on S7P *unc-22(RNAi)* and *β-cat-1(RNAi)* P-fragments at 14 dpc confirms regeneration of head identity.** *opsin+tyrosinase*: eyes (red arrowheads), *syt-1*: brain (red arrowhead) and ventral nerve cords (yellow arrowheads), *sFRP-1*: anterior margin (red arrowhead) and pharynx (yellow arrowhead), *fz4*: tail margin (yellow arrowhead). Scale bars = 250 µm.

Across metazoans, a gradient of canonical Wnt signaling activity is critical for establishing and maintaining the AP axis (reviewed in ^86^). In adult *Smed*, head regeneration is preceded by and necessitates inhibition of Wnt activity at anterior-facing wounds^54^. To test whether Wnt inhibition could elicit head regeneration in S7 P-fragments, we performed RNAi knock-down of *Spol-β-catenin-1* (*β-cat-1*) on amputated S7 embryos (**Figure 6B**). S7 embryos were bisected and soaked in *β-cat-1* or *unc-22* control dsRNA for 2 days, then transferred to Holtfreter’s media and allowed to regenerate until 14 dpc. Strikingly, *β-cat-1(RNAi)* resulted in head formation in 97% (82/84) of S7 P-fragments compared to 0% (0/85) of control *unc-22(RNAi)* fragments (**Figure 6D-F**). Regenerated heads in *β-cat-1(RNAi)* animals formed eyes (**Figure 6F**, *opsin + tyrosinase*), a brain (**Figure 6F**, *syt1-1*), and expressed markers of anterior identity (**Figure 6F**, *sFRP-1*). Some variation in patterning was seen, with fragments often regenerating fewer or more than two eyes (**Figure S4G**). *β-cat-1(RNAi)* S7 P-fragments exhibited normal gliding locomotion (**Movie 3**), indicating functional regeneration of the head. We confirmed efficacy and specificity of the *β-cat-1* knock-down by quantitative RT-PCR on pooled S7 P-fragments harvested at 14 dpc. RNAi treatment resulted in a significant decrease in *β-cat-1* transcript levels (**Figure 6C, S4B**), while relative expression levels of the paralog *β-cat-2* were unchanged (**Figure S4C**). Knockdown results were confirmed using two non-overlapping dsRNA constructs (**Figure S4A-H**). *β-cat-1(RNAi)* S7 A-fragments regenerated ectopic heads at posterior facing wounds instead of tail tissue (**Figure S4E-F, H-I**), phenocopying the *Smed-β-cat-1* knock-down reported in asexual adult regenerating fragments^90–92^. Altogether, these results reveal that S7 P-fragments are competent to undergo head regeneration in an altered signaling context and suggest that the ability to reset the main body axis after amputation is a rate-limiting factor for the establishment of whole-body regeneration competence.

## DISCUSSION

Construction of the adult body plan during embryogenesis and its re-establishment during whole-body regeneration are both dramatic and dynamic transformations. Differences may be readily observed in these ever-changing systems, but discerning necessary and critical regulators of these processes has proven to be challenging. Here we lay the foundations for rigorous comparative studies of molecular mechanisms regulating embryonic and regenerative development in a directly developing, long-lived animal with sustained postnatal regenerative abilities. We determined when *Spol* embryos were capable of mounting regenerative responses and discovered factors that were necessary and rate-limiting for the establishment of whole-body regeneration competence (**Figure 7A-B**). Our reductive approach focuses attention on the dynamic interplay between two critical components, stem-like cells and cellular purveyors of patterning information, in conferring regeneration competence. Our findings are consistent with proposed models of planarian regeneration^35,93^ and they speak to concepts and pathways that are broadly important for regeneration competence across species^94^.

**FIGURE 7:**
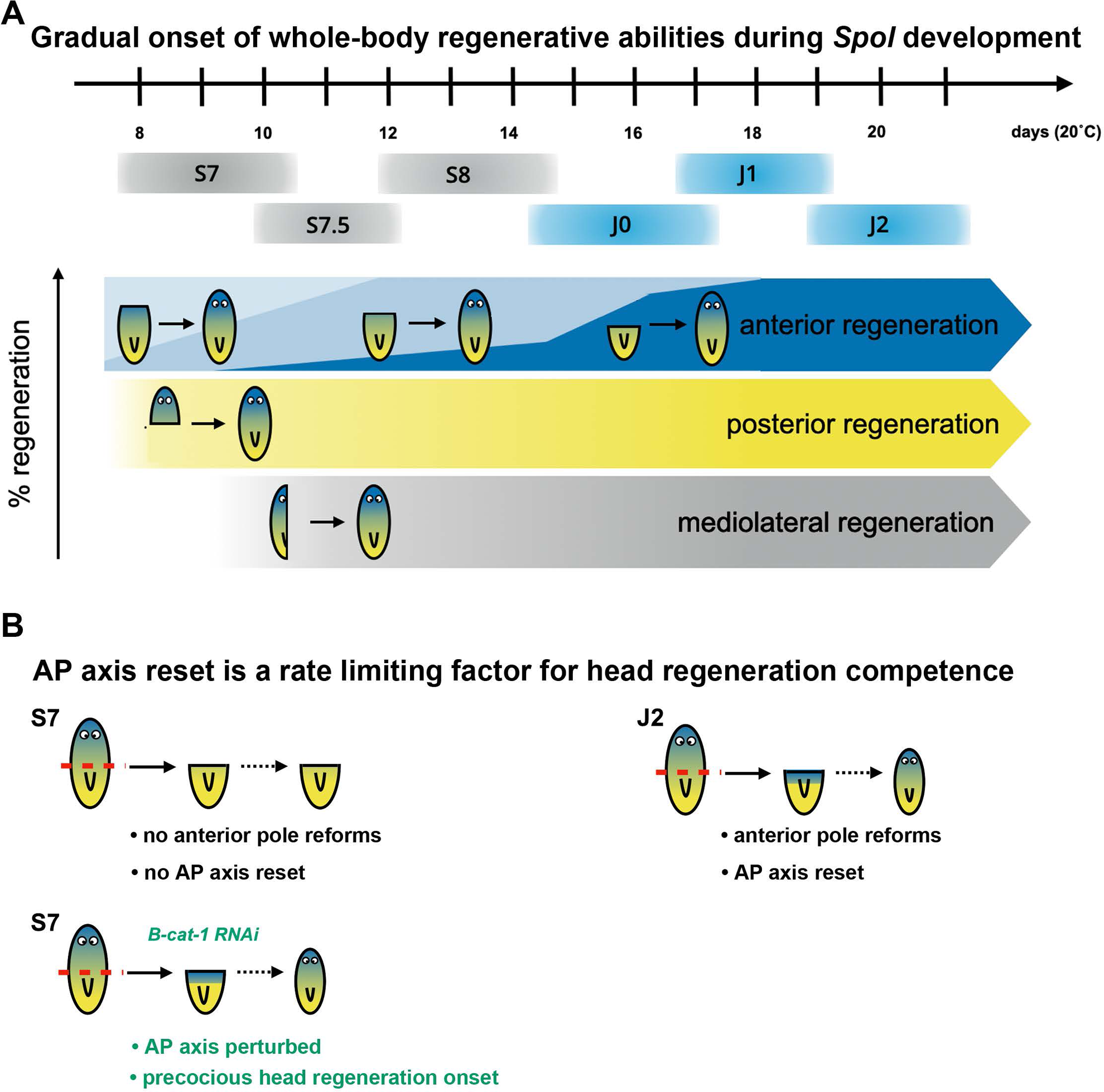
Whole-body regeneration competence is acquired gradually during *Spol* development and requires stem-like cells and robust AP axis reset mechanisms. **7A.** Whole-body regenerative abilities are acquired gradually during late embryonic and early juvenile *Spol* development. Onset of head regeneration abilities depends on developmental stage and axial position of the cut plane. In contrast, posterior and mediolateral regeneration abilities are constitutive from the earliest stages assayed (S7 and S7.5, respectively). We demonstrated that irradiation-sensitive cells are required, but not sufficient for regeneration of missing head tissue. **7B**. Main body axis reset is critical for whole-body regeneration. Headless S7 P-fragments do not re-specify an anterior organizer required for repatterning the AP axis. Remarkably, knock-down of the posterior determinant *β-catenin-1* following amputation is sufficient to induce precocious head regeneration in a developmental context that is normally head regeneration-incompetent.

### Stem-like cells and AP axis reset are required for whole-body regeneration competence

We report that irradiation-sensitive cells are required for regeneration in S7 embryos (**Figure 5 E-F**), that cycling cells are present through 48 hpc (**Figure S3C**), and that *piwi-1+* and differentiating epidermal progenitor cells persist in head regeneration-incompetent S7 P-fragments (**Figure 5G**). Head regeneration incompetence in late embryonic stages cannot be explained by *piwi-1+* cell loss, nor by complete blocks in *piwi-1+* cell proliferation or differentiation. It is possible that subtle axial- or stage-dependent differences in *piwi-1+* cell potency or activity contribute to observed head regeneration defects in *Spol* embryos. Although irradiation-mediated colony-forming assays have not been performed on *Spol* embryos, we would not expect *piwi-1+* cell potency to vary along the AP axis during development. In adult *Smed, piwi-1+* cells capable of clonogenic expansion and multilineage differentiation have been observed along the entire main body axis^95^. Next, we considered whether *piwi-1+* cells vary in their ability to respond to external patterning cues as a function of their location or developmental stage. We ruled out the possibility that embryonic *piwi-1+* cells are wholly refractory to cues required for regeneration because we observed robust posterior and sagittal regeneration abilities at the earliest stages assayed (S7 and S7.5, respectively), when AP axis reset was not required (**Figure 3B, I-K**).

We found that headless S7 P-fragments fail to reset the AP axis following injury: the anterior pole, a signaling center required to induce anterior patterning, is not reformed (**Figure 6A**). Strikingly, manipulating canonical Wnt signaling activity, an evolutionarily conserved regulator of AP patterning, via knockdown of *β-cat-1* resulted in precocious head formation in S7 P-fragments (**Figure 6B-F, S4**). In a low Wnt signaling environment, *piwi-1+* cells residing in S7 P-fragments produced a variety of anterior fated cell types (**Figure 6F**). Therefore, the repertoire of cell lineages produced by *piwi-1+* cells in S7 P-fragments is influenced by β-catenin-1 activity. This finding is consistent with previous reports of β-catenin-1-dependent expression of conserved Wnt target genes, like *sp5* and *abdBa,* in irradiation-sensitive cells from adult *Smed*^96,97^.

We speculate that failure to produce anterior pole progenitor cells^57–59^, a cell type necessary to implement the AP axis reset decision whose induction requires *β-catenin-1* function during regeneration initiation^54^, underlies the head regeneration defect in S7-S8 embryos. Failure to produce anterior pole progenitor cells may result from a *piwi-1+* cell-intrinsic defect impacting receipt of polarity cues during regeneration initiation. Alternatively, the primary cause of the defect may be within wound-responsive tissue(s) required for communication of the polarity decision to *piwi-1+* cells, such as the body wall musculature. We observed an “anterior first” bias in head regeneration abilities during S7-S8: anterior-facing wounds stood a better chance of regenerating heads when located at more anterior positions when AP axis reset was required (**Figure 3C-H, S2A-C**). It follows that anatomical system(s) required for instilling head regeneration competence may also display an anterior bias in their construction and/or maturation. In planarian embryos, the nervous system and body wall musculature develop progressively, with anterior structures starting to form before posterior structures, resulting in a gradient of structural completeness and maturation along the main body axis^73,98^. Changes in the composition of muscle subtypes and fiber density as the musculature forms may be reflected by changes in functionality of the tissue during development. In adult planarians, differences in muscle cell maturity have been reported to impact muscle fiber polarity and expression of axial patterning determinants following amputation^56,99^. Future investigations into position- and stage-dependent differences impacting regeneration initiation will help us to discern the root cause of this anterior regeneration defect. We favor the hypothesis that development and maturation of the body wall musculature dictates onset of position-independent head regeneration abilities, but do not exclude the potential necessity or involvement of other anatomical entities.

Finally, we consider whether precocious onset of head regeneration abilities elicited by *β-cat-1* knockdown could be due to activities other than AP axis reset. In many organisms, β-catenin functions in both the Wnt signaling pathway and as a component of the Cadherin complex, which is involved in cell-cell adhesion^100^. However, in *Smed*, the axial patterning and adhesion functions of β-catenin are split between two paralogs, β-catenin-1 and β-catenin-2. Smed-Β-catenin-1 is required for AP patterning^90–92^ but does not interact with the cell adhesion molecule ɑ-catenin^101^. In contrast, Smed-β-catenin-2 is not required for axial patterning^90,92^ but is capable of interacting with both E-Cadherin and ɑ-catenin^101^. Subfunctionalization of Spol-β-catenin-1 and Spol-β-catenin-2 paralogs, which are 98% and 94% identical to homologous *Smed* proteins, seems likely.

Canonical Wnt signaling regulates cell cycle progression, where it has been shown to license cells in G1 for proliferation via inhibition of GSK3 activity and promotion of *c-myc* transcription^102,103^. β-catenin-independent functions for Wnt signaling components have been shown to regulate mitotic spindle assembly, asymmetric cell division, and G2/M progression^104,105^. In adult *Smed,* β-catenin-1 may function in a genotoxic stress-triggered checkpoint that negatively regulates proliferation: *β-catenin-1* knockdown boosted mitotic cell numbers and induced head regeneration rescue in *Smed* adult P-fragments exposed to DNA damage^106^. Whether *β-catenin-1*-dependent changes in *piwi-1+* cell proliferation are associated with the precocious onset of head regeneration abilities in *Spol* S7 P-fragments is unknown. Wnt signaling interacts with other conserved signaling pathways and antagonistic, spatially opposed gradients of anterior-promoting ERK and posterior-promoting β-catenin-1 signaling have been proposed to exert pleiotropic effects on axial patterning, stem cell proliferation, and cell fate specification along the AP axis in adult planarians^87,97,107–109^. In future studies, we will determine whether stage- or position-dependent differences in ERK activation impact onset of head regeneration abilities in developing *Spol* embryos.

### Access to embryonic developmental programs fuels lifelong regeneration competence

In planarians, shared lineage specification programs executed in cycling, mesenchymal *piwi-1+* cells and their descendants are postulated to underlie the construction of the adult anatomy during embryogenesis, the maintenance of the adult body plan during tissue homeostasis, and the creation of new organs during regeneration^73,110^. Consistent with this idea, heterogeneous expression of fate specifying transcription factors for prominent developmental lineages spanning three germ layers was first observed among the cycling *Smed-piwi-1+* population at the outset of organogenesis during S5, and persisted in all subsequent life cycle stages^73^. Organogenesis programs during embryogenesis and postnatal regeneration may proceed using shared transcriptional modules to direct differentiation of required cell types, though the activating signals and sources of progenitor cells are likely to be distinct. Vetting of this hypothesis will require demonstration of functional necessity for fate specifying transcription factors during adult anatomy establishment, as well as rigorous comparative studies of transcriptional regulatory strategies used during embryonic and regenerative development. Studies in other highly regenerative organisms also support the idea that regeneration re-uses many embryonic gene regulatory networks, albeit with some differences in the wiring of these circuits^111–115^. Redeployment of embryonic patterning programs may occur in some, but not all vertebrate regeneration paradigms. For example, limb regeneration in the *Xenopus* tadpole requires formation of a specialized signaling center within the wound epidermis that directs expansion and differentiation of underlying mesenchymal progenitor cells^116^, likely through redeployment of the limb outgrowth program used during development^117^. In contrast, axolotl tail development and regeneration appear to proceed using separate strategies^118^.

Planarians likely sustain the activities of axial polarity pathways that specify the definitive body axes during development throughout postnatal life^119^. Maintenance of embryonic axial patterning organizers and constitutive polarity signaling activity throughout adulthood enables stem cell-dependent transformations in axial identity during homeostasis^90–92,120,121^, and occurs in some other whole-body regeneration-competent animals, like *Hydra*^122^. Planarian AP patterning pathway effectors may promote transcription of chromatin modifying agents required for transcriptional plasticity in differentiated tissues that mediate early phases of the amputation response upstream of polarity reset^123^. In contrast, animals that lose regeneration competence as they mature may undergo progressive silencing of injury-responsive enhancers required for transcription of critical regeneration genes, like *wingless* and *Wnt6* in *Drosophila* imaginal discs^124,125^, or *Shh* in the *Xenopus* limb^126^.

### Mechanisms impacting changes in regeneration competence during development may help to explain changes in regeneration competence across species

Absence of head regeneration in clades with robust posterior regeneration abilities is a phenomenon seen in other Spiralians, including Annelids, Nemerteans, and Platyhelminthes^127,128^. Comparative phylogenomic and regeneration studies suggest that head regeneration abilities were acquired once or more within the Triclad Order: some planarians undergo axial position-independent whole-body regeneration, some exhibit graded head regeneration abilities along the AP axis, and some lack regenerative abilities entirely^66^. The progressive acquisition of head regeneration abilities during *Spol* development will likely hold true for closely related species^129^ and mirrors some of the variation in head regenerative abilities described among adult planarians^66^. Knockdown of *β-catenin-1* induces precocious onset of head regeneration abilities during *Spol* development (**Figure 6, S4**) and rescues anterior regeneration in head regeneration-deficient species^66,87–89^. These results strongly support the hypothesis that axis reset is a critical point of regulation for whole-body regeneration competence and underscore the pivotal role played by canonical Wnt signaling in determining the outcome of the AP polarity decision. Importantly, these results also demonstrate that changes in regeneration competence may be evoked by changes in the expression or activity of a single gene product. Considering studies of regeneration competence across both life cycle stage and evolutionary context together can help identify critical conserved regulators of regenerative ability, which may be fruitful therapeutic targets for regenerative medicine.

## MATERIALS AND METHODS

### Animal strains, husbandry, and developmental staging

*S. polychroa* (3n = 12) hermaphrodites (Nico K. Michiels, Alejandro Sánchez Alvarado) normally reproduce asexually through sperm-dependent parthenogenesis^68–70^. In asexual embryo production, sperm is required for oocyte activation and embryo development, but the paternal genome is excluded from the offspring^69,71,72^. Animals were maintained in 1x Montjuic water ^130^ at 20°C in constant darkness and were fed twice weekly with pureed grass-fed beef liver paste. Egg capsules were collected daily. Following collection, egg capsules were disinfected by soaking in 10% bleach for 3 minutes, washed liberally in 1x Montjuic water, and stored in dated polypropylene containers at 20°C in constant darkness. The egg capsule collection date is considered 1 day post-egg capsule deposition (dped). To maintain optimal fertility and fecundity of the breeding colony, sexually mature adults were replaced every 6-7 months with juveniles. *Spol* reach sexual maturity within 6-8 weeks post-hatch provided they are fed 1-2 times per week.

Staging series modifications: S6 was subdivided into S6 (5-7 dped spherical embryos) and S6.5 (7-8 dped elongating embryos). S7 was subdivided into S7 (8-10 dped) and S7.5 (10-12 dped). S7 embryos are elongated, with small visible eyes and definitive pharynx primordia. Pigment cells, if present, are sparse and restricted to the dorsal side. Primary gut branches are visible, with few secondary branches. When positioned ventral side up, S7 embryos cannot curl their heads, nor can they twist to right themselves. Onset of gliding locomotion occurs during S7. S7.5 embryos have better defined gut branches, particularly in the anterior half of the embryo. When positioned ventral side up, S7.5 embryos can curl their heads, and they can twist to right themselves. S7.5 embryos are motile.

### Amputation assays

Amputations were performed with microknives (Fine Science Tools #10316-14). S7 and S7.5 embryos were cut and cultured in 1x Holtfreter’s media^73^ supplemented with 100 µg/mL gentamicin sulfate (Gemini Biosciences #400-100P); fragments were gradually acclimated to 1x Montjuic + 100 µg/mL gentamicin sulfate beginning at 2-3 days post-cut. S8 embryos, J0, J1, and J2 juveniles were cut and cultured in 1x Montjuic + 100 µg/mL gentamicin sulfate.

### Plasmid DNA constructs

*Spol* gene fragments were cloned into pT4P^131^, pCR™Blunt II-TOPO™ (ThermoFisher 450245), or pDL1^132^ plasmids. pDL1 was a gift from Daniel Lobo (Addgene plasmid # 182263; http://n2t.net/addgene:182263; RRID:Addgene_182263). Primer and confirmed insert sequences for each construct can be found in **Supplementary Table 1**.

### Whole mount in situ hybridization

Colorimetric WISH was performed as described^133–134^ with the following modifications: Intact J2, S7 A-, S7 P-, J2 A- and J2P-fragments at 14 dpc were transferred to clean polystyrene petri dishes, rinsed at least six times in 1x Montjuic water, and were immediately treated with 5% N-acetyl-cysteine (NAC) in 1x PBS for 6 minutes to kill the animals and remove mucus. 10,000 Rad-treated embryos (7 dpi) and S7A and S7P 3 dpc fragments were treated for 4 minutes with 5% NAC in 1x PBS. NAC treatment was omitted for intact S7 embryos (unirradiated or 10,000 Rad-treated, 3 dpi) and unirradiated S7 A- and P-fragments fixed within 24 hpc. Fixation was carried out in 4% formaldehyde in 1x PBS-0.5% Triton X-100 (PBSTx) for 1 hour at room temperature, with occasional swirling to ensure animals did not stick to the dishes or to one another. Fixed embryos were washed for 2 x 5 minutes with PBSTx and underwent gradual dehydration into methanol (5 minutes x 50% methanol, 2 x 5 minutes in 100% methanol) and were stored at −20°C until use. Following rehydration, S7.5 and older embryos/fragments were bleached in 1% formamide, 6% H2O2 in 1x PBS-0.1% Tween (PBSTw) for 30 - 60 minutes under bright light. Proteinase K treatment and post-fixation steps were omitted. Colorimetric WISH samples were imaged with a Zeiss AxioZoom v16 microscope and Axiocam 208 color camera.

### Immunostaining

Immunostaining was performed after fluorescent WISH development^133,134^ using DyLight 488 (ThermoFisher 46402). Samples were incubated overnight at 4°C in anti-H3S10p (Millipore 05-817R-I) 1:1000 in WISH blocking buffer. Secondary antibody incubation was performed using anti-Rabbit Alexa 555 1:1000 (Abcam 150086) and DAPI 1:2000 in PBSTw + 5% Normal Goat Serum for 2 hours at room temperature. After post-fixation for 20 minutes in 4% formaldehyde in PBSTw, samples were cleared by incubation in 10% Triton-X for 48 hours at 37°C followed by Ce3D+ overnight at room temperature^135,136^. Samples were mounted between coverslips in Ce3D+ and imaged on a Nikon Crest X-Light V3 spinning disk microscope. Maximum confocal projections were created in Fiji/Image J.

### Irradiation

7 dped egg capsules were soaked in 10% bleach for three minutes, rinsed liberally in 1x Montjuic water, and divided into two treatment groups: unirradiated controls and irradiated egg capsules. 10,000 Rad irradiations were performed in 15 mL falcon tubes in 1x Montjuic water + 100 µg/mL gentamicin sulfate on a rotating stage in a Mark 1 Cesium Irradiator (J.L. Shepherd & Associates, San Fernando, CA, USA). Irradiated egg capsules were transferred to 10 cm petri dishes following the irradiation and water exchanges were performed every 1-2 days thereafter.

### RNAi

dsRNA was synthesized from HT115 bacteria transformed with pDL1 plasmids^132^ harboring 400-600 bp insertions for *Spol* transcripts of interest. *unc-22* dsRNA was used as a control^137^. dsRNA synthesis and purification from bacterial pellets was performed as described in^138^. S7 animals were cut immediately prior to soaking in 10 ng/ul dsRNA in 1x Holfreter’s + 100 µg/ml gentamicin for 48 hr. After 48 hr, animals were washed into new dishes with 1x Holfreter’s + 100 µg/ml gentamicin and gradually transitioned into 1x Montjuic + 100 µg/ml gentamicin over the next two days. Fragments were monitored for 14 dpc.

### Quantitative real-time PCR (qPCR)

Total RNA was extracted from pools of five fragments per RNAi condition at 14 dpc using Trizol (Invitrogen 15596018). cDNA was synthesized from 162.5 ng total RNA using the SuperScript III Reverse Transcriptase kit (Invitrogen 18080093). qPCR reactions were prepared using iTaq™ Universal SYBR® Green Supermix (Bio-Rad 1725125) and were run on a QuantStudio™ 5 Real-Time PCR Instrument (ThermoFisher A28133). Data were analyzed using the 2^−ΔΔCT^ method^139^, normalized to the housekeeping gene *Spol-EF2* as described in^140^. qPCR primer sequences are found in **Supplementary Table 2**.

## ACKNOWLEDGEMENTS

We thank members of the Davies lab and the NCI-Frederick Cancer and Developmental Biology Laboratory for feedback and discussions of the work, and Dr. Alejandro Sánchez Alvarado for his foundational support of planarian embryogenesis research and gift of the *Spol* triploid strain used herein. This Research was supported by the Center for Cancer Research, National Cancer Institute, National Institutes of Health Intramural Research Program project number ZIA BC 0120009 (ELD). The content of this publication does not necessarily reflect the views or policies of the Department of Health and Human Services, nor does mention of trade names, commercial products or organizations imply endorsement by the US Government.

## AUTHOR CONTRIBUTIONS

Conceptualization: CLTB, BCS, CAS, ELD. Methodology: CLTB, BCS, CAS, ELD. Validation: CLTB, BCS, CAS, NTK, ELD. Formal Analysis: CLTB, ELD. Investigation: CLTB, BCS, CAS, NTK, ELD. Resources: ELD. Writing-Original Draft: CLTB, ELD. Writing-Review and Editing: CLTB, ELD. Visualization: CLTB, ELD. Supervision: ELD. Project Administration: CLTB, ELD. Funding Acquisition: ELD.

## SUPPLEMENTAL INFORMATION

Supplemental Figures 1-4 and associated Figure legends.

Table S1.

Table S2.

Movies 1-3, accompanying Figures 2 and 6.

