## Supplementary Online Materials for "Axis reset is rate limiting for onset of whole-body regenerative abilities during planarian development"

**SUPPLEMENTAL FIGURE 1** – ***(associated with Figure 2)***

**S1A.** Representative live images of S7 - J2 animals.

**S1B.** Whole mount in situ hybridization (WISH) on S7 and J2 animals show developing structures and body plan organization across stages. *sFRP-1:* anterior margin (red arrowhead) and pharynx (red asterisk). *syt1-1:* nervous system, including the brain (red arrowhead) and ventral nerve cords (red arrows). *opsin* + *tyrosinase*: developing eyes (red arrowhead). *foxA1:* pharynx (red asterisk). *fz4*: posterior region. *wnt-1:* posterior pole (black arrow).

**S1A-B:** Anterior: up. Live images: dorsal views. WISH images: D: dorsal. V: ventral. Red asterisk: pharynx. Scale bars: 500 µm.

**S1C.** Distribution of *Spol* spontaneous hatches as a function of time post egg capsule deposition (dped) at 20˚C. Approximately 80% of hatches occur between 15-18 dped. 98% of gravid egg capsules hatched within 30 dped. n= 189 hatched egg capsules.

**S1D:** Percent surviving Anterior (A-) and Posterior (P-) fragments at 14 days post-cut (dpc) as a function of developmental stage (S7, S7.5, S8, J0, J1, J2). S7 A (n=46), S7.5 A (n=52), S8 A (n=53), J0 A (n=58), J1 A (n=42), J2 A (n=39). S7 P (n=49), S7.5 P (n=52), S8 P (n=53), J0 P (n=58), J1 P (n=42), J2 P (n=39). Data points: replicate means for three independent experiments. Error bars: standard error of the mean.

**S1E:** Percent P-fragments that regenerated 2 normally patterned eyes (dark gray), >1 eye (white), or no eyes (maroon) by 14 dpc at developmental stages S7 - J2. S7 P (n=31), S7.5 P (n=51), S8 P (n=49), J0 (n=58), J1 (n=60), J2 (n=56).

**SUPPLEMENTAL FIGURE 2** - ***(associated with Figure 3)***

**S2A. Left: P1 fragments regenerate normally patterned eyes at high frequency**. Percent P1 fragments that regenerated 2 normally patterned eyes (dark gray), cyclopic eyes (light gray), 1 eye (white), no eyes (maroon) by 14-16 dpc. **Right: Eye regeneration quality for P1 fragments improves during development.** Fraction of regenerated eyes that were mispatterned in P1 fragments as a function of developmental stage. One way ANOVA, ($F(3,11)=15.26$, $p=0.0003$). Tukey's multiple comparison tests: S7 vs S7.5: p=0.0024. S7 vs S8: padj=0.0005. S7 vs J2: padj=0.0005. S7 P1 (n=36), S7.5 P1 (n=48), S8 P1 (n=48), J2 (n=64), S7 P1: 3 independent experiments; S7.5, S8, J2 P1: 4 independent experiments. Error bars: standard error of the mean.

**S2B. Left: Incidence and quality of regenerated eyes in P2 fragments increases during development.** Percent P2 fragments that regenerated 2 normally patterned eyes (dark gray), cyclopic eyes (light gray), 1 eye (white), no eyes (maroon) by 14-16 dpc. **Right: Eye regeneration quality for P2 fragments improves during development.** Fraction of regenerated eyes that were mispatterned in P2 fragments as a function of developmental stage. One way ANOVA, ($F(3,11)=7.628$, $p=0.0049$). Tukey's multiple comparison tests: S7 vs S7.5: ns. S7 vs S8: ns. S7 vs J2: padj=0.0123. S7 P2 (n=51), S7.5 P2 (n=59), S8 P2 (n=60), J2 P2 (n=69), S7 P2: 3 independent experiments; S7.5, S8, J2 P2: 4 independent experiments. Error bars: standard error of the mean.

**S2C. Left: Incidence of regenerated eyes in P3 fragments increases during development.** Percent P3 fragments that regenerated 2 normally patterned eyes (dark gray), cyclopic eyes (light gray), 1 eye (white), no eyes (maroon) by 14-16 dpc. Right: Fraction of regenerated eyes that were mispatterned in P3 fragments as a function of developmental stage. S7 P3 (n=46), S7.5 P3 (n=62), S8 P3 (n=62), J2 P3(n=77), S7 P3: 3 independent experiments; S7.5, S8, J2 P3: 4 independent experiments. Error bars: standard error of the mean.

**S2D. Percent surviving sagittal L (gray) and R (white) fragments at 21 dpc as a function of developmental stage (S7.5, S8, J0, J1, J2).** S7.5 L (n=45), S7.5 R (n=45), S8 L (n=45), S8 R (n=45) J0 L (n=46), J0 R (n=46), J1 L (n=45), J1 R (n=45), J2 L (n=45), J2 R (n=45). Data points: replicate means for three independent experiments. Error bars: standard error of the mean.

**S2E. Percent sagittal fragments at 21 dpc that regenerated the missing eye as a function of developmental stage (S7.5, S8, J0, J1, J2).** A modest increase in eye regeneration incidence was observed as development proceeded. One way ANOVA, ($F(4,10)=5.482$, $p=0.0134$). Tukey's multiple comparison tests: S7.5 vs S8: ns. S7.5 vs J0: ns. S7.5 vs J1: p=0.0247. S7.5 vs J2: p=0.0117. S7.5 L+R (n=57), S8 L+R (n=82), J0 L+R (n=92), J1 L+R (n=90), J2 L+R (n=88). Data points: replicate means for three independent experiments. Error bars: standard error of the mean.

**<u>SUPPLEMENTAL FIGURE 3</u>** ***(associated with Figure 5)***

**S3A.** WISH on intact S7 (top) and J2 animals with riboprobes against the cycling cell marker *piwi-1* (left), the early post-mitotic epidermal progeny marker *prog-1* (middle), and the late post-mitotic epidermal progeny marker *AGAT-1* (right). Anterior: up. Live images: dorsal views. WISH images: D: dorsal. V: ventral. Scale bars: 500 μm.

**S3B.** Percent surviving unirradiated and 10,000 Rad treated S7 A- and S7 P- fragments at 7 dpc (10 dpi). Four independent experiments. Unirradiated S7 A: n=134. 10,000 Rad S7 A: n=162. Unirradiated S7 P: n=132. 10,000 Rad S7 P: n=162.

**S3C: Mitotic cells persist in headless S7 P-fragments.** Maximal confocal projections of anti-H3S10p staining on unirradiated S7 A- and J2 A-fragments (left) and S7 P- and J2 P-fragments (right) at 48 hpc. Anterior: up. Scale bar: 250 μm.

**<u>SUPPLEMENTAL FIGURE 4</u>** ***(associated with Figure 6)***

**S4A.** Schematic showing the *Spol β-catenin-1* transcript and locations of two non-overlapping dsRNA constructs used for knockdown and the qPCR amplicon. All Figure 6 knock-down data was collected from experiments using dsRNA 1.

**S4B.** RT-qPCR measurement of *β-cat-1* levels in A- and P-fragments at 14 dpc. Both *β-cat-1* dsRNA constructs produced significant decreases in expression compared to control RNAi (adjusted p-values: 0.0161, 0.0213, 0.0036, 0.0032; Šídák's multiple comparisons test). Error bars: standard error of the mean. Four independent experiments.

**S4C.** RT-qPCR measurement of *β-cat-2* levels in A- and P-fragments at 14 dpc. No significant difference in expression of *β-cat-2* was measured in any knockdown condition compared to control (Šídák's multiple comparisons test). Error bars: standard error of the mean. Four independent experiments.

**S4D.** Percent of *unc-22(RNAi)* and *β-cat-1(RNAi)* S7 P-fragments at 14 dpc for both knockdown conditions that regenerated heads. Error bars: standard error of the mean. Four independent experiments.

**S4E.** Experimental schematic for knockdowns. S7 animals were cut and soaked in 10 ng/ul *unc-22* (control) or *β-cat-1* dsRNA (dsRNA 1 or 2) for two days and scored for regeneration at 14 dpc.

**S4F.** Percent of *unc-22(RNAi)* and *β-cat-1(RNAi)* S7 A-fragments at 14 dpc for both knockdown conditions that regenerated posterior-facing heads. Error bars: standard error of the mean. Four independent experiments.

**S4G.** Percent of *unc-22(RNAi)* and *β-cat-1(RNAi)* S7 P-fragments at 14 dpc that regenerated heads with 0 eyes (gray), 1 eye (orange), 2 eyes (blue), or more than 2 eyes (green). Error bars: standard error of the mean. Four independent experiments.

**S4H.** Representative live images of *unc-22(RNAi)* and *β-cat-1(RNAi)* (dsRNA 1 and dsRNA 2) S7A- and S7 P-fragments at 2dpc and 14 dpc. *β-cat-1(RNAi)* S7P-fragments regenerate heads with eyes (red arrowheads) and S7A-fragments regenerate posterior-facing heads with eyes (red arrowheads). Scale bars = 250 μm.

**S4I.** WISH on *unc-22(RNAi)* and *β-cat-1(RNAi)* (dsRNA 1) 14 dpc head fragments with anterior biomarkers to demonstrate posterior-facing head regeneration. *β-cat-1(RNAi)* A-fragments regenerate anterior structures at the posterior wound site and have reduced expression of posterior markers. *opsin+tyrosinase*: eyes (red arrowheads), *syt-1*: brain (red arrowhead) and ventral nerve cord (yellow arrowhead), *sFRP-1*: anterior margin (red arrowhead) and mouth (yellow arrowhead), *fz4*: tail margin (yellow arrowhead). Scale bars = 250 μm.

**MOVIE 1** (*Associated with Figure 2).*

Representative *Spol* S7 A- and S7 P-fragment regenerates at 14 dpc.

**MOVIE 2** (*Associated with Figure 2).*

Representative *Spol* J2 A- and J2 P-fragment regenerates at 14 dpc.

**MOVIE 3** (*Associated with Figure 6).*

Representative *unc-22* and *B-cat-1 RNAi* soaked *Spol* S7 A-fragments at 14 dpc

**SUPPLEMENTARY TABLE 1**

Primer and insert sequences for plasmid DNA constructs used to generate riboprobe templates and dsRNA.

**SUPPLEMENTARY TABLE 2**

Primer sequences used for RT-qPCR experiments.

### Supplemental Figure 1

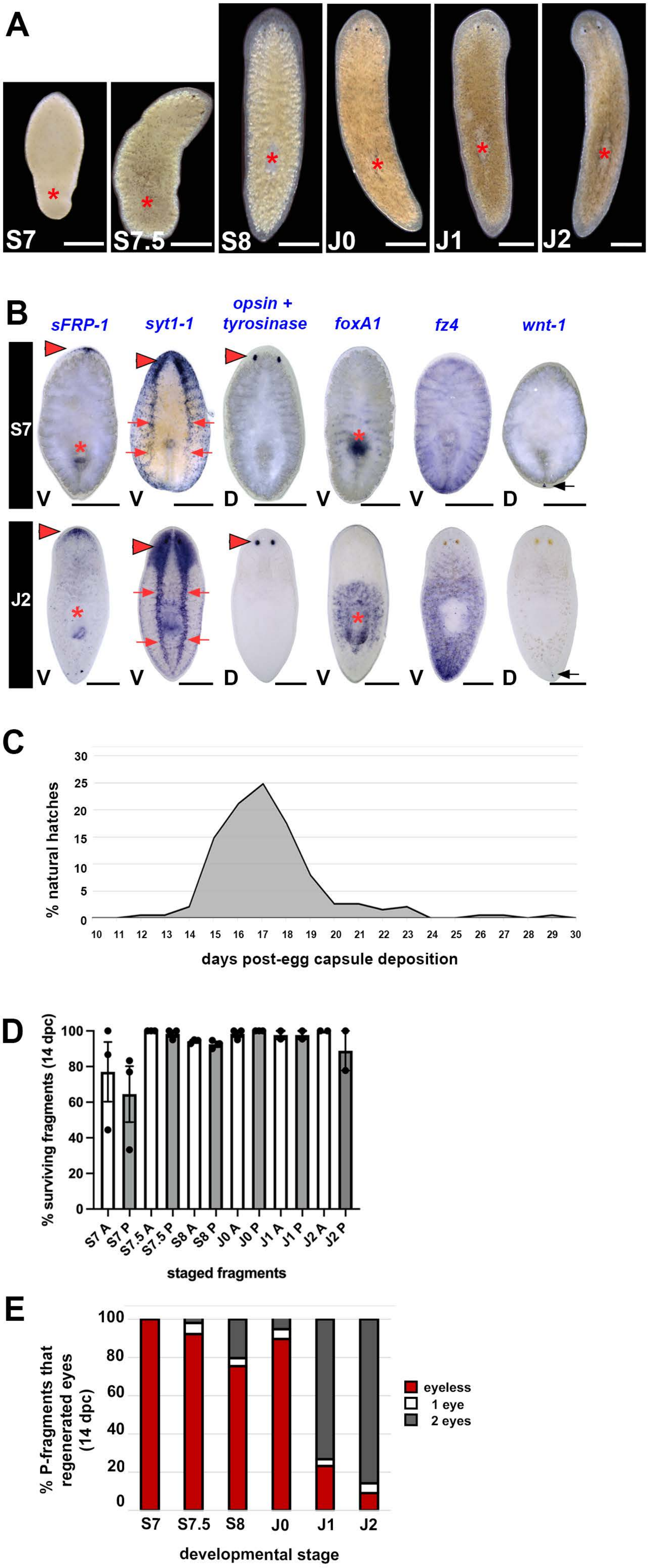

### Supplemental Figure 2

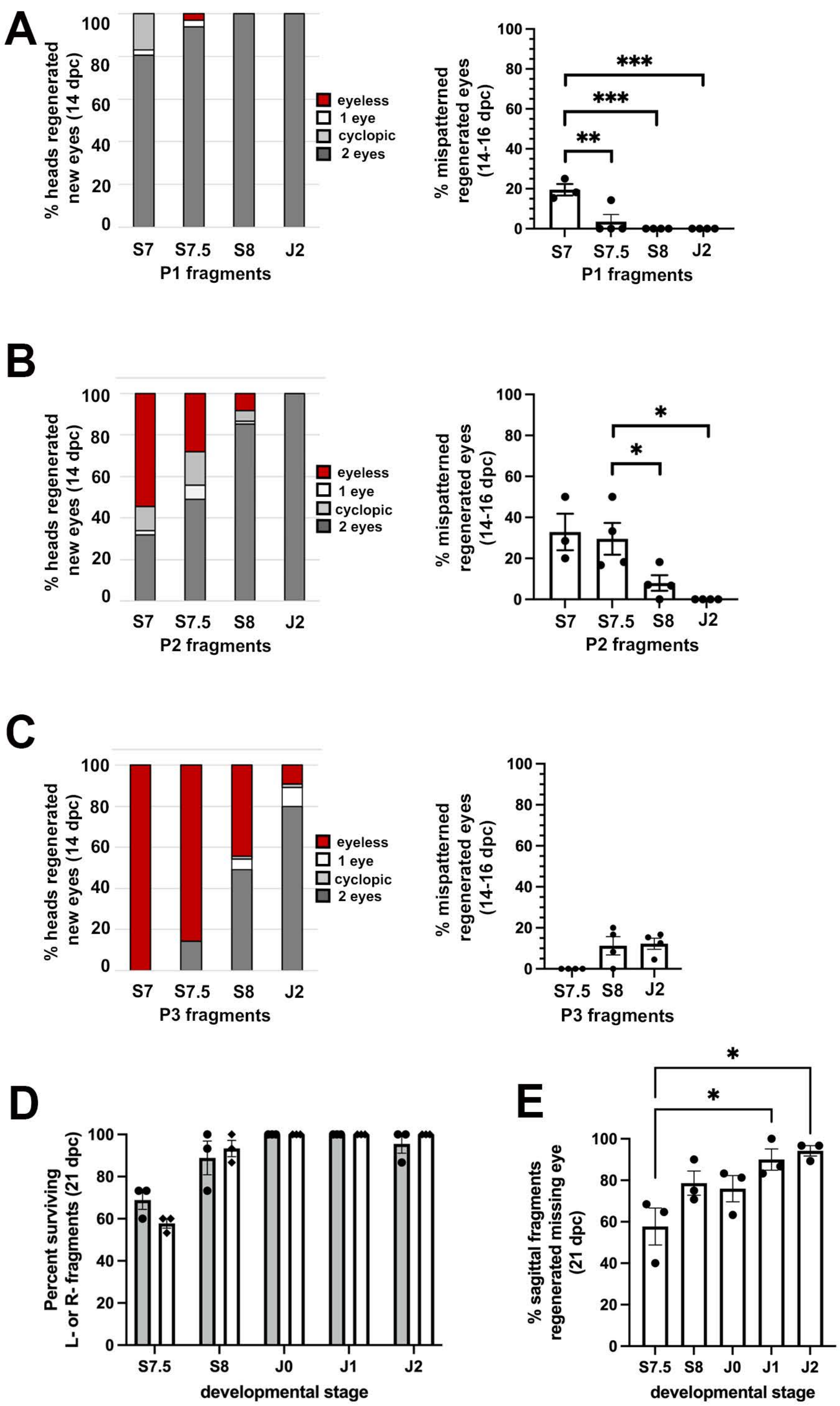

### Supplemental Figure 3

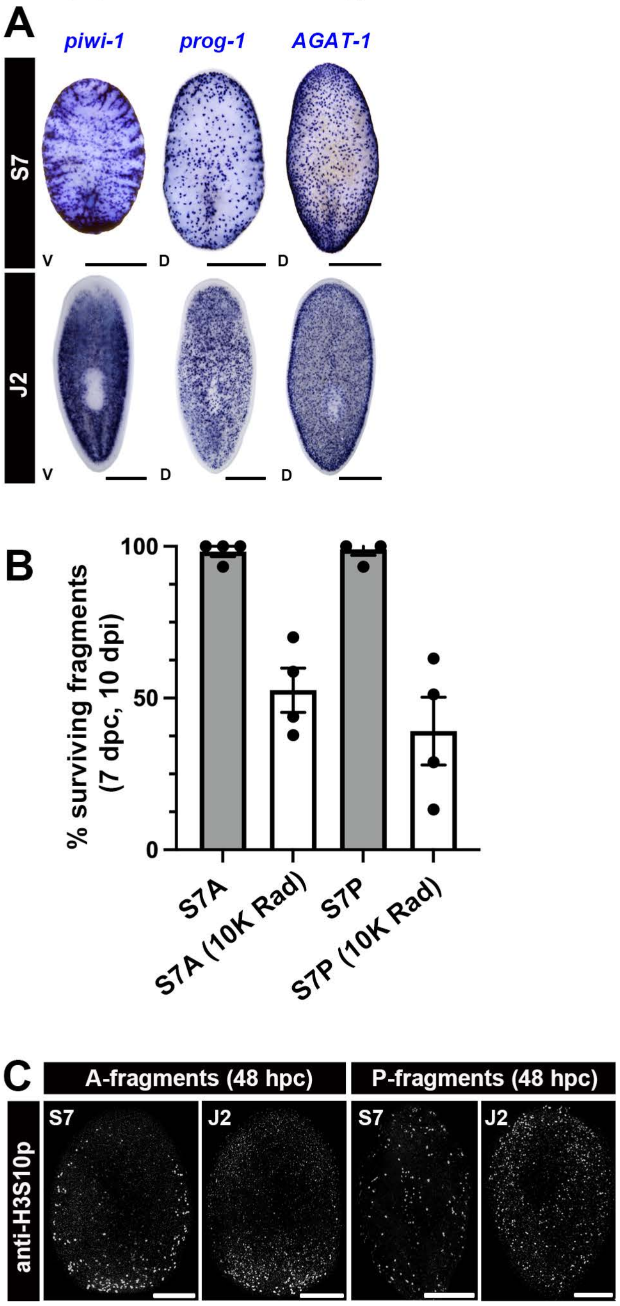

### SUPPLEMENTAL FIGURE 4

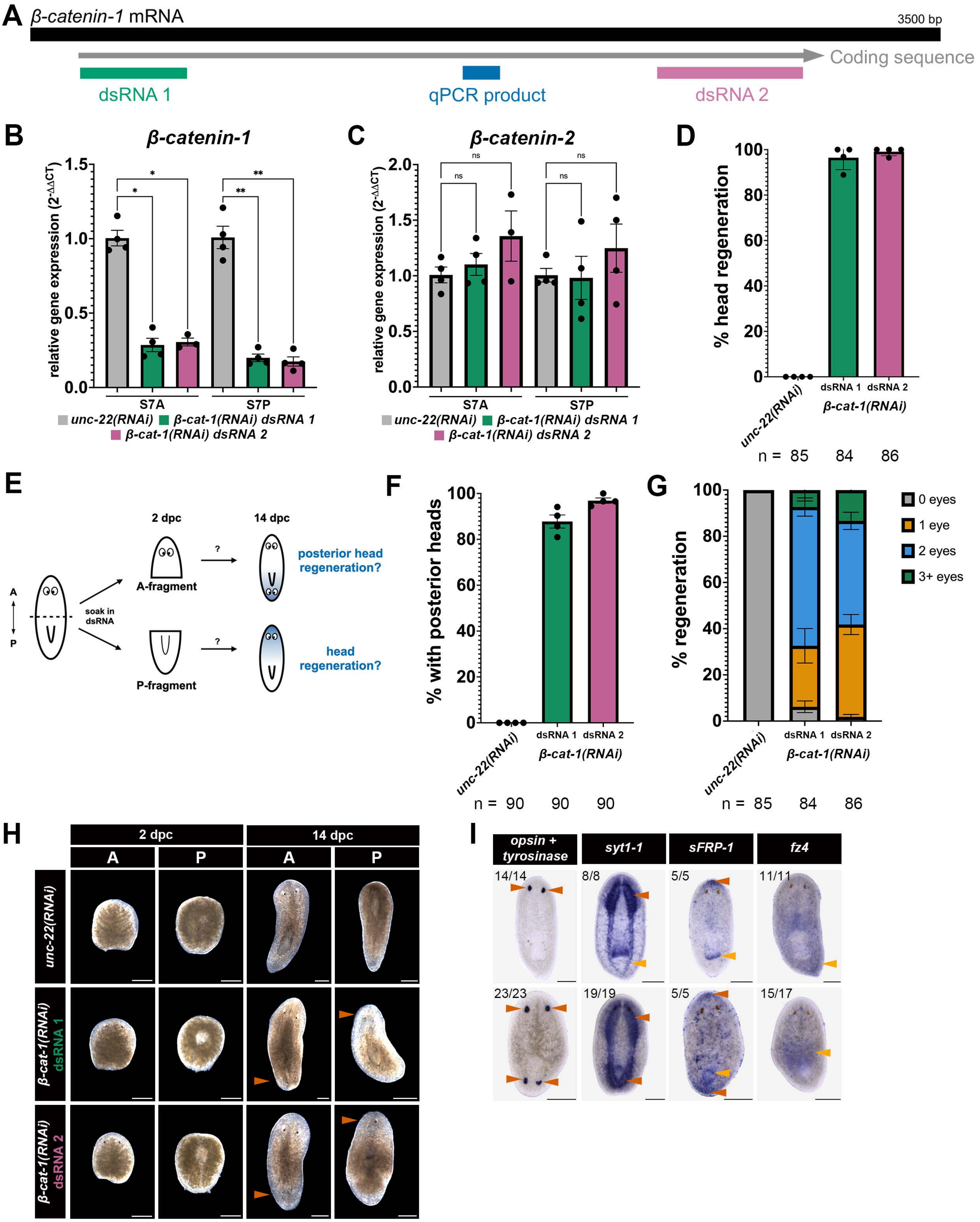

Supplementary Table 1

| Construct | Fwd primer | Rev primer | Vector | Insert length (bp) | Insert sequence |
|---|---|---|---|---|---|
| *Spol-B-cat-1* dsRNA 1 | tcgtcaattctcctcaagcag | tctggtatcgcactttcagc | pDL1 | 410 | tcgtcaattctcctcaagcagatgaatttataactgacaaaacaaatttcacaagaacatggcagcaaaatcaatatcttatcgattctggaataaattcagctttagaaagtcatagtcacagtgtcagcagcaaacatggatatgatgatatggattctgaagaccaaaataaaaatcttaatgaatggaaatatccttcaaataccaatacaaatgatggcttgtctaatgatatatgtatgctaactccagatactcctggtagtacaagttgtttaagacatggcggtgaaggtatggctgatgtaagtttgtcggcttgttacgataataaagaagaaaataacagaaatcctgattttggaattgatatagataaaatagaagctgaaagtgcgataccagat |
| *Spol-B-cat-1* dsRNA 2 | acgatatgtagctcctccgc | ccagtccgcatccaacatac | pDL1 | 565 | ccagtccgcatccaacatacatacatccatcatatcctttttagaattttcatttgacgaacacaaatttccactaatcgataaagaatgagatgatgggctcttgaaactggaatttgaataagtcttttgaacatcttgattaacacattcattgttaagatttatagagcccatctgtttgaagtgtctcattccgatttcagaattcccagtttgatttatcaaattgtaatttggatctacgtaatttggattgtacggatatcgattttgcggttgtagagtattagaaaacgtcatgtggtaatctgtaacatccacatcattgattggacataaagtttcatttgtagcattattttgatcaccacattgcgttactgaattgtgtctatggctgacacatggcgattgattgtgcataatcggagaattcaaaatcgaatttaaatatccagaattattggcttgattagctggtaaagatatcgatagattgtttcttagattattcgggttttgaggaatgatattgtagttgtgcggaggagctacatatcgt |
| *Spol-syt1-1* | caatgagaggggctattacgc | ggcgagcatgaaaacgaatg | TOPO | 1427 | ggcgagcatgaaaacgaatgtggaaaaattttggaatgatctcggaaaaaaattacacatggatccaaaagcactaataggagtaacagtcggtttgggaatattgtttctgctgttcctttattgtttatgtaaaaaatgtgttttcaagagacgaaagaagaaggaaactgcgaagaaaggagccaaggggggtagtcgatatgaaaagtgcccaattgctgggaaactcatataaagagaaaattcaacccgatcttgaagaacttgatgcaaacatggaagacaatgaaggtgtaaaggaaaaagaagacgttcatttaggaaaactacaatattctttagattatgactttcagaaaggagaattaactgttggagttattcaagcaactgatcttccagctatggacatgtcaggaacatcagatccatatgtcaagttataccttctcccagataagaaaaagaaatttgaaacaaaggttcataggaaaattttaaatcctgtattcaatgaaacatttgttttcaaagttcctttcaatgaagttgcatcgaaaactttgattttcaatgtttatgactttgacagattttccaaacacgatcaaattggtcaaataaaagtaccactgggagctatagatttaggtcgagtaattgaagaatggaaggaactagaatctccagaaaacgatggggaaaaggaaaacaggttaggagacatttgtttctctttgcgatatgtaccgacttctggaaagcttaccattgttattttggaggcaaagaatcttaaaaaaatggatgtcggaggattgtctgatccatatgtgaaactctccttgatgcttaatggtaaacgagtgaagaaaaagaaaacaacaatcaaaaaatatacactaaatccttactacaatgaatcattttcatttgaagtaccattcgaacaaatacaaaaagtaaacttgattgtcaccgttgttgattatgacagaattggaaccagtgaaccgattggtcgaattgttctaggatgtaacgcaacaggagcagaattgcgacattggtcggacatgttggcgaatccccggcgaccaatcgcccagtggcatacccttcaagagatgccagaaaataattgattgtgctcatttgagcggtttgtattcttcgtgaaataagttatcaaaatttgagaaatgggtttttttcctgatataccagagttcatttgtgtgtggtttcgttaaatattttggtggtgttgcatcttatcgtttgtcctctaatctaataatgttttattctttatatagctgatttattct |

| | | | | | |
|---|---|---|---|---|---|
| | | | | | cgagcgaaacatttttatgtgtaccaaaatacttacttatgtcatagccattgtgattagtctgtaaactattgcgtaatagcccctctcattg |
| *Spol-opsin* | tatggcattgggaccctgaa | cattcgtatgtccactcgtga | TOPO | 1155 | tatggcattgggaccctgaatttgaatcaattgtccatccgtactggcggacttttgatatggtgccggaagtttatcattatttagttggtgtgtatatttctattgttggaatatcaggggttttgggaaatctccttgttctttacatatttgcaagcgccaaaagtttaagaactcctccaaatatgtttataatgagtttagcaattggggatttgaccttttctgctgtaaatggctttcctttactcactatatcaagttttaacactcgatgggcttggggaaaattgacgtgtgagatctacggtttcatcggtggtctttttgggttcatatctatcaacacaatggcactgatttctctggatagatattttgttattgcccaaccatttcaaacaatgaaatcactgacaatcaaacgcgcaataattatgttggttttcgtatggctctattctttgatttggtcgacaccgccattctttggatatggaaattacgtcccagaaggatttcaaacgtcttgtacattcgattatttgacccaatcaaagggtaacataattttcaatattggaatgtacataggaaatttcataattcccgttggaataataatattttgttactatcagattgtcaaagcagttcgagtacatgaattagaaatgctaaaaatggctcaaaagatgaatgcgtctcatccaacttccatgaaaaccggcgcaaaaaaggctgatgttcaagctgcaaagatttctattataattgtctttttatatatgttatcatggacaccatacgcaataatagctttaatggctcttactggacgaagagatcatctgaatccatacactgcagaattgccggtactctttgctaagacctcagccatgtacaacccatttatctacgcaataaatcatccaaaatttagaattcagttggaaaagaaatttccctgtttgatttgttgttgtccaccaaaaccaaaagaaagggacactacaagtgcagcatcagcaacaaaaccaagtaaatatcagagcgaaaaagaaagttcgatgtctcaattggatactgaagacgataaacaagtcgtagaaacgaccaagaatcaaatcacgagtggacatacgaatg |
| *Spol-tyrosinase* | tgatccagatacaaatgacgtga | ggtctttcctccttctcctca | TOPO | 534 | tgatccagatacaaatgacgtgagaatgaaagacataggaaaaggactgacccatcaaaatgtcatgattcataatatggttcatagttttattaatggaacttggatggatttcactacgagtcccaatgatccattatttttactgcaccacacatttatggacaaaattttcgaaatttggcttctgagaaatacactatctaaatacccggagaaattttcaataccaattggacagcatagtgatgactttattgtgcctttcataccattaatcagaaataagaatttgtttactccggcaaataagttcggctatgcttacgataacaatattattggaaataaattattcacaaacgacccactagatttcgaaaagagctcgtcggaaatttactacattctaacggcggttgtcattgtagttgtagttgccacccttgtagggctgtattgcttcagacctaaccaaataagtcacacgacttataaagtcattgaagaagatgaggagaaggaggaaagacc |
| *Spol-notum* | gcgaatactcctttagcctgc | acaataatacctgtgaatcccgt | pT4P | 751 | gcgaatactcctttagcctgcctcaaagtctgagctgcaaaaccacctaattgttgaatgtatgaaaactttttgttagatcctccttccaaaatcggaactttagacatttgtagttgagcgtcatcaaataaactttgtataatgaaaactggatttttcaaatgtctatatatgacagggcccataaaacatttccattctctaccgcgaccttggaatttttttgcaccttcttggaatccgaggattccataatttcattccctttcttaaagcattttctggaggacactcgtaaatatgagtacagtttgactgcctatacgctggataatcgagaaaccatgctgaatcgactattccatgaataaatactttccgtttaagtttttttcttcaatcgtcttttcaatctatctatattcaacaaaactccaattcctccggcgctcgatccaacaaaagctacttcttgaaccttttttaaaatgctgattttgggtcaaatcatcaacgactgtgtccaaaattcgagatccatgaaaatataatccattagtgttttctaattgtttaccagaccacaaatcactcgaacagtacggaatatacactgaattgtaatcatggaaattaggattgattcgttcattattagaaagaattccaccaagttgccgctcagaagaccaaaattctgatgaaaataagtcataatgggtacgtctacgggattcacaggtattattgt |

| | | | | | |
|---|---|---|---|---|---|
| *Spol-sFRP-1* | tcggaactttctcgtttgttgt | agactttgctttcctgacatttt | pDL1 | 1315 | agactttgctttcctgacattttattcaacatatggattccgtgtaacgtccatatgtaacggcccgaatgatcagaataaattggatgagtgacacagtcaattactttatttttagcagtatgtgggcaatttgttttgtttggttgaagttttcccatcattttaatcaatatcatgcaataaagtagaggaaaatgaggctgtattcagcatgtgaaaatttaacattgtgacatctgacggtaattaatgtgattggctagtttcaacttattagacacaaaacacaagctattaactgctataatatttaatcaaatgccccatatgaaatgaattcatggaaatgaccacgatttttcccgtaagtttattattgttcatctccaatgtccatagtttcatcggagattggcaatcatttcaaagcggttacacaattgatcaatgttatcaaattcctgataactttaccctttgtagtaatgtcggttacatgcgaatggttttacccaattatattcaacatgagtctattgaagaggctgctcagcattctaaagtgtggatgggtttagttaacacacaatgccatgccgatataaaaaagtttatttgttcactttatgctccggtatgcataagcaatcagagaattcaaaaagttccgccttgcagagaactatgtgtaactgttaaggaatcatgtttgccaagcatgaaaatgtatggatttgactggccaacaattatgaaatgttcgaaatttccacaatttaaaaattcattatgtatacctaaaaccgttgtgccaggtaaaaagtgtgacctatgcatgaaaattgcgtcttatgaagtcatagcgaacagattctgtttatctccgatagtaatcagagcaaaaatcaaaagaattattccaaccaacggcaatgccattcagattatcctacaaaagaaatcaaaattcctcaaatttcaagccaacttcagtcagaacgctcttgagttggagctgaaatgcaactgtacaaatctgaaattcaaacaacgcgccttgaaaggacgctggattattatggcccaaattgacaggggcaataaagctgtggtgaattttatttccaaatggaaaagaaagaaagcagaattcaagcattcaatgaaaattattcagaactttggtcatattgtttgtaaatcaaaactgccggacggaatcagtttacgggaaagatatgctgcctatttaaaaatgaacaatcaattaaaatattcaaatgaaagaaaacaacaaacgagaaagttccga |
| *Spol-fz4* | tgccgaatttagttggaagc | ttggtttggatcggttcttc | pT4P | 1576 | ttggtttggatcggttcttcctctttttcctttcccagaatcgaagcgttgaaatgcaagacttaacggtttttttgttgccccatatccaaataccggaagtgattccaataactaaattcatgaaaatcctcaacatacgcatctcaactgatgggaggaccggctgtaaaatttcaatgcatttcatgcttgaccagtttatcccattgcggccaatacaattcggtgaaaaccgaaacaatgctttgagtgtgatatcaaaatttgttaatttgtacgcatcaattatgtaactgacgacggtgcacaaaacaggcaagacataaagaattgaaaaaaccccaattttagccattaacttttcaagacgctttaaattgttggtgtcaatacaatccggtcgctgcaaaaacgaacgacgaacctttaacaaggaaacatatcccatagttagaaacacaattcctgttaacaaataaatcatgtttggtacaataacaaaaccccaaagaactttgttgttactgttcccaacaaagcattgtcccgtcaactcgtcagcgtcgattttgtggagaataagaataaaaatggacttcaaagctggaatagcccaactcaacaaatggagcatactgctgatggattcaatggcttcgtagccccaatgtcgagaagctgaaagaaaccaagtaatcgtcagtacgacccaccataaggcactggcggaaccgaaaaagtaaaccatcaaaaagttaatcttacaccaagtggtctctctaccggaactgacaatgaaatcggttttatcatcaaaggttcgacaagcaaccacatttcgacccaaaatcgccccaaacagatttccactagcatagaaaaaataacagatagaaagaaaaacaatcggcttttccggataaagaaaccgattcgtgttgcaggagaatacgatgatagtcatcaaacaggaaaagaaacaaagtactgaccaaccgagcagccaaatattggcaaagactttatcatgcgcattgaactggacgtttgccgaacagatttgagcgcatgaaatattcggagtttgcccaatgttgacctctgacggcaagcattgcgttcccaccattttgctcattttccgataaatgtcgagttttgattgatcaacgcgatttccttgtataatgagcaggagttttctataaaggtcttgatagctttcctcttgactatttgtcgacggatgaaattcgtcctgggaaaacgaatccggagttaagcagatttttccccgttctgggaatttttcaca |

| | | | | | |
|---|---|---|---|---|---|
| | | | | | gtccaattcttctgaccaggataacttgaatgtttctttcatcaacggttcacatcgttctttgacatgattgcataaatgtttgcacggtttcactgtcgacgatgacatcgaggtgtctgtttcacacatcggaaaatagacagagcaaagaaaaagtttcaaataattgctgcatttgatgttgatcaacggattgaaatctctgaaccgtttgcttgcttcgatctgtgaatagcttccaactaaattcggca |
| *Spol-follistatin* | gcaatatagcgagttctacgaca | agaatccacacagttcaacgag | pDL1 | 349 | aagaatccacacagttcaacgaggccattttgcattgagaagacattgtccagaatgaagaacttcaataaaaacattattttgacatgataaaaaaatctttatagcaggtatttctcgtttcgtagttagttgacaaacacaacaattccacagttttataaatgccattctcacattgacgcattgatgtgggacatttgatacaaactggtgatccattcctatggtgagtaatgcacgtcatgtcactttttttgacattcgacattcatacaatttgcgtcagttctgcaaactccactataggcaacctgaatttctttctctcgtagaactcgctatattgc |
| *Spol-foxD* | cactattcggtcagcgctct | tacatggccaaagtcgaagc | TOPO | 1137 | tacatggccaaagtcgaagcgtttgtggtatgaactgattagcataagattcttcagattgtttcgctttattatctgttgaatcatctgaaattatgtgagaaatggaaaatttctgaaatgtagatagttcgtcttctgacgaattgagtttacatcttttgtttactggttcgttgtattcagtttttgatgttgatgggataactattctcccatcatgtggataagtggaactaacgatccaatgaactcatgccctctgaaattattctcaaatggccgcggatatcgaatagcattcaacgggggaatgatgttattcgggttgggaataattgatttggtaatattttgatgaaataacaactgatttacgaggttttgctggactgtcattggattaggtggaatagtcattctgaatgaagctggaggaattatcaaatgagattgattgtgattaaacatttctgacggcaattgacgtttataccgttttcgtctacgtaaaaaactaccattatcaaacatatcttctgatcttggatctaatgtccaataatttccttttccaggatttcccggttctcttggtatttttatgaagcaatcatttaaagataaattatgacgaatactattttgccatgctggaaatctatctttataatatgggaaacgtcccattataaactcacaaattccactcaatgttaattttctttgaggagatcttaaaattgccatagtaataagagctatatatgaatatggaggtttaacattatgagatttagatttacatcgatcttgagacatcggaattttattatcagtttctttttttgtcttctgaacatattcgatctaaatcgctttctgaatgtatattatcatcactttcgcattctttcaagtcatctacatccacatcatcatctcgaagtaaattttttatttgttcaattgtctgagaatctttgaatgcctgaggcatatcttgttcatatttcatgtttattctctgagacgctcttaaatattcaatagaatttgatctgtgatatggcaataagtaagagtcatctccgaaattcttcattaattttgcatcagagcgctgaccgaatagtg |
| *Spol-wnt-1* | acattcgactttgcaacacca | accgatttacccgctaacag | TOPO | 928 | gacattcgactttgcaacaccatatgaatttgcaattgcaattttccattattgttttagatcttgttagaaaagttcggttgcagcaaagattattgcaactgtcagtagaattcgacaaggaattacatatccgtcccgtcgtgcccagaatgtgccattcgggttttgaatcacaataaaagtttttttggaacatcttctaaataaactaaatctttagcatatggatattgtttttctatttgaatgaaaatatgtttctttgctattatgagtgctttgcgatattgttgatttttattagtagattttcttttctgcgaatcggcgagtaagttaacttcactatcttctttggcgaccttgacggcattttcatatttttctttcaaaattttaccgatttcgtcaatactagcaacttttctgtggcaaattttagtcgtacaagaaccacttgtgccctggcagacacatttcacgttcatacttttgactgcaactcttctacctgcccccgtgttgtgaaggttcattaatgtctttttgttccaatctggactgtctggatcaaacattttcctcgcaaatctccgtccaaattgcacattgtcatcacatccttgccaaatccagttggtttgagatattcgtcccttattattgcacggacagtgtgataacttgaacgagcaagcttcggcgacagtttgagcgacacttgcactaagcatcgcataaatgaaagcggtttcgggaaatcctttgagcataatgtcaccaaaaagtaaggcggatggattattcagattcggtgtcggacagttccagcgatgattggcaaa |

| | | | | | |
|---|---|---|---|---|---|
| | | | | | caacttttgacatgtgtaaatacccttctaattccttcaatagccacttggattagatcgctgttagcgggtaaat<br>cggt |
| *Spol-foxA1* | acttttccgaacatgcg<br>cat | gtcactgttgcctggttg<br>ag | TOPO | 1845 | acttttccgaacatgcgcatgcagcctgcaaattatttcaaaataagccaatcaaattgaagaaaaattcaa<br>gagaatataaattaacttcgcgaatagactcaaaatttgtattggagcaatgtgtgtgcctcattgtgagtggtt<br>gcgatcgcatctcaaattatttacacacagaaaataacgcacacatacattgaatagatatatttgatcaagt<br>caatatcacaatgtcaatattagctttttctgaagaaagaagacgaactagaagaaaaacaacaacgaaat<br>agattcgataaggctactgagcgatttttataatctaaatattattttattattgatacggacaaaaattggtcttta<br>ccaagtactactagattgttgataaagagatgcttggaaaaaatccctatgaaactgcaatgagcaacgtgt<br>attctctacctccaggaggttctatttacaatatgaatccaatgagtatatcctcagctggctacaactctcagc<br>aagtatcaacactatcattgaacttgactggaattggacctcattcattaagcccaatgagtgcaagtatgtct<br>ggcatagctgcaatggccggtggaatgagacaaggtcttgaattaggtcttggtagaagtgatagtccgag<br>agataaaaattctatttccaataacaaccgaccatatcaaagaagttacactcatgccaagcctccatacag<br>ttatataagtttgataacaatggcgattcaaaattcgccagtaaacatgtgcactctatctgagatctatcaattc<br>attatggatcattttccatactatcgtcaaaatcaacagcgatggcagaattcgattcgacactctttgtcctttaa<br>tgattgttttgttaaggttagtagaagtccggaaaaaccaggtaagggctcatattggacactacatcctcaat<br>caggaaacatgtttgaaaacggttgttatctcagaagacaaaagcgattcaaagatccacaccgagaaatt<br>ggtagacagagtcaaagagctgccactggtccaggatcaaacatcacagaaaacaatcatgacaatgca<br>tcacaagaagctagtgataacgctgaaagtgatacgaaacccaacatcaaacaacttgatttatcaaatga<br>tctcttaactaatcaaggtcataatattaaaaatactaatccaacttctgttagtcagagttgttcgatgtttcatcg<br>gaaaaaggaaaactgctcaccagtagaaatgaaattgaataatcaaacccaacaaccaaaccagcaa<br>gaacatccacaaattcattacaatcccaatcagcaattctactcaaatcagcaaaacattttccaacaaagtt<br>ctttagatcactatagcttattggcatcagatgaccctcttggtcagggtatgcacttgccaccaggtgcgaata<br>gtgttttcggactttatggaggacacaacttaccaaacgatgatcaaatttctgtgtcattaccatcgatatcatt<br>atcgggacatccgtatgacaatttatcaacagctatggcatatcaatacgaagcatctcaacacaattcgtca<br>ttactaacaacaagtaatccgttctcaatagatcgtttgatgcatccgagactagttgctgccgccatgggtgtg<br>agtccccatgatactctatacgcaggagccactggctcatcagttgatctcgaacacatgaaatactactcaa<br>actacaacaacgtgcctccctatgcatctgcaatgtctgactactacaaatatgtacaaaatcctcaaccagg<br>caacagtgac |
| *Spol-piwi-1* | atggatgcaacgaatg<br>ttacta | taagcgttcacgaattct<br>gtc | pT4P | 2415 | atggatgcaacgaatgttactagtgatccaaatttgagaccaagaagaggtggtctccaaagacgattcgc<br>aataatagaacctgccttacagccagacacagttacagataaaattggtaaagatggacgagtagttagtct<br>caaatccaactacgctaactttaacattaaagtcgatgggccattttacatgtacgatgtagtaatggaaaata<br>agagtattaggactaagcttgagttattattagcgtatgctaacagtaaaaataacaaccaaccattttgttttg<br>atggtagaagattgtttacgacttccaagtggcattcaggcgaagatgctgaaaatttggtgatcaacaatgtt<br>gaaatttccataagattgttaacaacatttttgaaaaattctgaagaatattatcaaatggttaatattgtatttaat<br>aatattcagatttttatgggtcaagaaaaaattggcagacaatttttctgggttcaaattgtgaaagtggtggca<br>ttcgagaaggacctagctttttaaaatggacaatttcaagattttaggagggtattctacaaccattacaagag |

| | | | | | |
|---|---|---|---|---|---|
| | | | | | gaacaaatgacaatggaactgcttcaaccttactctatcttgaaagaatcaaccgcgttcttaatgaaaacagtgtagtatcagtgtatcgacgagatcaaatcgatcaattaataggaagagatatcataacgaaatataataacaagacgtatagaatttccgacattaaagaaatggatgtgaacgacgaagttcaattgggcgacaagacgatatcatatttaaattactttaaacagcgttacaatattaatttgaagaatgacaagcagccatttgttatatctagggtcaaacgttcgatggttcgtccaaaagaaaaaaccgaaaatgagcccgaagggcctaaagaaacagatcaaagtttatcgattcctggtgaattatgctttttgtgtggattttcggatagtgaaagaagtaatataaatttgcaaaagaatttaggaagtgtattgaaacgtgaacctagagaacgattgaatgatattagagatttttgtaaatggacaggagctggtaaatcaaaagattatatgactagttggggtatgagtattgaagaacaaccaattacaattcgtggtcgtgagttggcaccagttgatatattttcaaatgaagtaaaaattaattctaaaattcccgatgattggaaatttggtgagattaaatttgatgttcctaaaggtacaactcatcgttttggtgtattggttgttgatagaaacccaagtcacttcaataatttcattgaagatgttaagagagaaattggaagattgagaataaattatacaatggacaatatttcgtcgtgtcgatctaatgaagttgaagatgctttagctgattttattcgtgttaataaagtccatttggcattagtattcattccagatgataaagtttatgctaaggtgaaaaatttcaccatgtctactggccttttgactcaatgtgtgacacagagaaatggtagcaatagagatgatagacgtcgtaaaaccgttgcagataaatctgttatgcaaattttctctaaattaggatatgatccttggggcatcaatctcaaaatggccccaactatgatagtgggattggatacttttcatagtaaaacaggaaagcggtctgtacaagcatcagtattttcaattagcgctaaattttcacaatatattagttttgtgaattcttcaaagggcaaaaatgaatttcacgaaaacttgggtaaaaactttctcactgcaattaccactttccaaaacaagttcaatactatgccccatcgcttgattatttatcgtgacggtgttggagactcacaattggcatttactaaaaaatttgaaactgatgctgttatgaagatgattgaaaagatttatgagaatcagactttgccgcaaataatttacgttgtagttaaaaagcgtattagcgttaaatttttcaaagatggcggtaatccaaatcctggtacggttgttgatgaaaaaatagtgaagccaaatttctatgaattttatttagtgtcacagaaaacaactaaaggaactgcaactccgacgaattataatgttcttatggacactaaatttacaaataagaagacaaatgaagtgtcagtgatgtcacctagtgtcttgcaacaaattacttattcgttaacgcatttgtattttaattggatgggcactattcgagttccagttccaacgcattatgcacatcgtttggctgaactcgttggaaagattcatcgtggcgttactcctccgactattaatgacagaattcgtgaacgctta |
| *Spol-wnt-1* | acattcgactttgcaacacca | accgatttacccgctaacag | TOPO | 927 | acattcgactttgcaacaccatatgaatttgcaattgcaattttccattattgttttagatcttgttagaaaagttcggttgcagcaaagattattgcaactgtcagtagaattcgacaaggaattacatatccgtcccgtcgtgcccagaatgtgccattcgggttttgaatcacaataaaagtttttggaacatcttctaaataaactaaatctttagcatatggatattgtttttctatttgaatgaaaatatgtttctttgctattatgagtgctttgcgatattgttgatttttattagtagattttcttttctgcgaatcggcgagtaagttaacttcactatcttctttggcgaccttgacggcattttcatatttttctttcaaaattttaccgatttcgtcaatactagcaacttttctgtggcaaattttagtcgtacaagaaccacttgtgccctggcagacacatttcacgttcatacttttgactgcaactcttctacctgcccccgtgttgtgaaggttcattaatgtctttttgttccaatctggactgtctggatcaaacattttcctcgcaaatctccgtccaaattgcacattgtcatcacatccttgccaaatccagttggtttgagatattcgtcccttattattgcacggacagtgtgataacttgaacgagcaagcttcggcgacagtttgagcgacacttgcactaagcatcgcataaatgaaagcggtttcgggaaatcctttgagcataatgtcaccaaaaagtaaggcggatggattattcagattcggtgtcggacagttccagcgatgattggcaaaca |

| | | | | | |
|---|---|---|---|---|---|
| | | | | | acttttgacatgtgtaaatacccttttctaattccttcaatagccacttggattagatcgctgttagcgggtaaatcggt |
| *Spol-prog-1* | gaatgccgtcctcaagaatt | tgtcccttttcacactgatc | pT4P | 554 | tgtcccttttcacactgatcaacttttttatattctccgtcacaatgcgcatccaacgcgtcgtattctgcgtcacaatggccatcaattgaatcataaacgcccataaaccaattttcaaaagctcgcaattctgctcgacatttttgtcatatttgctgccacattttgaatgataaccttcccatctttttgagtaatcattcaatgcattcacatatctacaatcacaatctttgtcgagcttcaaataaagggggtcgcaatatttatcagtttcttcgtataatttcatgcaatctggtttcgtgacctttcccttcaatttctcttttaagctttctccttcctcttcttccaattctttcactggcttctttcgtctacttgatttagcgggagcttttccagaaaccttggaatcagtctcttcctcatctttatcagaacctttatccgaaacttttccatctttcgagttatcagcagcctcctcttccagcttagattttactctcctcatgaagtccaataattcttgaggacggcattc |
| *Spol-AGAT-1* | atttccaccggttttctgtg | tcctcgccttctaatatccg | pT4P | 1210 | atttccaccggttttctgtgatataaattgtacaaaaatgcttaccaaaatcgctagatccctgcacgatcagatatctataaatgttatcaataaattgtcggctaggttatgctcatatacaccgggtaacaaaaaacacagcccagtttgggcctggaatgaatgggaccccttagaagaaattgttttgggacttccagacatggcatgttgtccagagattttaccagaagccaaagcttgcatttcagaaaacaagttcgattttcttttaaaacataaaggaaaatattggaaggatgttattgaaccagaacattacaaactgatgttagatgagcacaagaattttataaagattttagaaggagaaggagtaaaagtcgtacacccagaaccgattgatttcggtaaaacactctgtactcctaacttttcggcccatggagtcgattgtgcgatgccgagggattttctcttaattgttggaaatgaaatgattgagtcaaccatgacttggagatcgagatatttcgaatttatgtgttataaaaaattaattaaagactactggaaccggggagccaaatggacagctgcaccaaagccaatgtgtaaagaaacactttttaaagaaaattatcgagcagacgaagattcaacggcatttttttgatgaaaagacttcaattctaactgaagaggaaccggttttcgatgccgcagatttcattcgatgtggtagggatatatttgcgcacgtcagtcaggttagcaacaacgctggaattgaatgggtacgcagacatttggctcccagaggaattagagttcattccatgaatttcgtagatcctaaacctatgcatatcgacacgacacttttcacgcctcggccaggtctcgcaatatcatgtccagaaagaccgtgtttacaaattggcatgtttaaaaaagccggctgggatgttgtcgttgctccatatccggatatttcgaaagattttccatattatattagctctcaatggttgtctcttaatgttgtcatgttggatgaaaagcgagttctcgtagaaaaggatgaaattggaattcaaaaattgtttgaaaagtgtggcatcactcctgtaaaggttcctttcaaacactgcttttctataggaggtggtattcattgctggacatcggatattagaaggcgagga |

Supplementary Table 2

| Transcript | Fwd primer | Rev primer |
|---|---|---|
| *Spol-β-catenin-1* | GACATTTTCGAGCCGTGCAT | ATCGGGACCCAGTTGCATTT |
| *Spol-β-catenin-2* | ATTGAGGACACAGACACCGC | TCTCCGAAGCAAGTCCCAAA |
| *Spol-EF2* | GCGAGCCAGAAGATTTGTAT | TGAATATCTCCCATAGGTCCA |
